# Core and accessory effectors of type VI secretion systems contribute differently to the intraspecific diversity of *Pseudomonas aeruginosa*

**DOI:** 10.1101/2022.04.11.487527

**Authors:** Antonia Habich, Alibek Galeev, Verónica Chaves Vargas, Olga Vogler, Melanie Ghoul, Sandra B. Andersen, Helle Krogh Johansen, Søren Molin, Ashleigh S. Griffin, Daniel Unterweger

## Abstract

Bacteria use type VI secretion systems (T6SSs) to deliver effector proteins into other cells or the extracellular space. Those effectors kill microbes^1^, manipulate eukaryotic cells^2^, and sequester nutrients^3^. Which T6SS-mediated functions are generalisable across bacteria of a species or are specific to particular strains is little known. Here, we use genomics to test for the intraspecific diversity of T6SS effectors in the opportunistic pathogen *Pseudomonas aeruginosa*. We found effectors that are omnipresent and conserved across strains acting as ‘core effectors’, while additional ‘accessory effectors’ vary. *In vitro* and *in vivo* experiments demonstrate different roles of the two types of effectors in bacterial killing and virulence. Further, effectors compose various effector combinations. Within one local population of clinical isolates, we observed 36 combinations among 52 bacterial lineages. These findings show the distinct contribution of T6SS effectors to strain-level variation of a bacterial pathogen and might reveal conserved targets for novel antibiotics.

## Introduction

Bacteria benefit from frequent interactions with their biotic and abiotic environment. By secreting effector proteins directly into target cells or the extracellular space, bacteria kill other microbes^1^, manipulate eukaryotic cells^2^ and take up nutrients^3^. In this regard, the type VI secretion system (T6SS) has been shown to be an effective protein delivery tool of numerous Gram-negative bacteria, including symbionts and pathogens^4–7^. Its function is mostly inferred from experimental work on a few reference strains of diverse species.

Generalising findings on the function of the T6SS in a particular bacterial species is not trivial. T6SS-mediated phenotypes are expected to be highly specific to the T6SS effectors of the respective strains. It is the effectors and their diverse enzymatic activities that mediate anti-prokaryotic, anti-eukaryotic and nutrient-acquiring activities, ultimately leading to phenotypes such as killing of prokaryotic and eukaryotic cells^8–11^. Experimental studies showed that effectors and their corresponding immunity proteins, which protect sister cells from getting killed, are often encoded side-by-side on mobile genetic elements and are subject to recombination^1,12,13^. Consequently, some effectors are known to vary between strains^14–17^, but most effector loci have not yet been systematically analysed in a bacterial population of one species. Although synergies between effectors were reported^18^, the combinations in which effectors occur remain mostly unknown. Inferring the function of the T6SS for a bacterial species is therefore close to impossible without knowing the intraspecific diversity of T6SS effectors.

We investigated the intraspecific diversity of *Pseudomonas aeruginosa* T6SS effectors, which are both a model system for the T6SS field and a virulence factor of this opportunistic pathogen. *P. aeruginosa* is the main driver of chronic lung infections in cystic fibrosis (CF) patients^19^. The detection of T6SS components in the sputum of CF patients and recent reports of T6SS-mediated colonisation resistance to *Burkholderia* in the CF lung demonstrate the clinical relevance of this secretion system^4,20,21^. Each T6SS effector is (i) translocated by one of the three types of T6SSs (H1, H2, and H3)^9,22–28^, (ii) associated with the secretion machinery by Hcp, VgrG or PAAR-domains^29–32^, and (iii) targeting nutrient uptake, prokaryotes and/or eukaryotic cells^1,9,14,25,33^. To our best knowledge, more T6SS effectors are known and characterised in depth for *P. aeruginosa* than for any other species. They are therefore ideal for systematically characterising the prevalence of effectors in a bacterial population and testing the impact of T6SS effector diversity on bacteria-bacteria interactions and pathogenicity.

Our analysis focused on 22 T6SS effector-encoding loci in the available genome sequences of 52 phylogenetically distinct *P. aeruginosa* isolates (herein referred to as 52 distinct clone types) collected from a local cohort of 33 CF patients^34–44^ (Fig. 1a, Supplementary Table 1). The large number of isolates in this collection and their extensive diversity uniquely enabled us to study the variation in T6SS effectors in one local population and later expand our analysis to isolates from various sources from all around the globe. What causes diverse *P. aeruginosa* bacteria to behave similarly in some aspects and not in others could be influenced by differences in the intraspecific diversity of T6SS effectors and subsequent variation in T6SS function.

**Figure 1.**
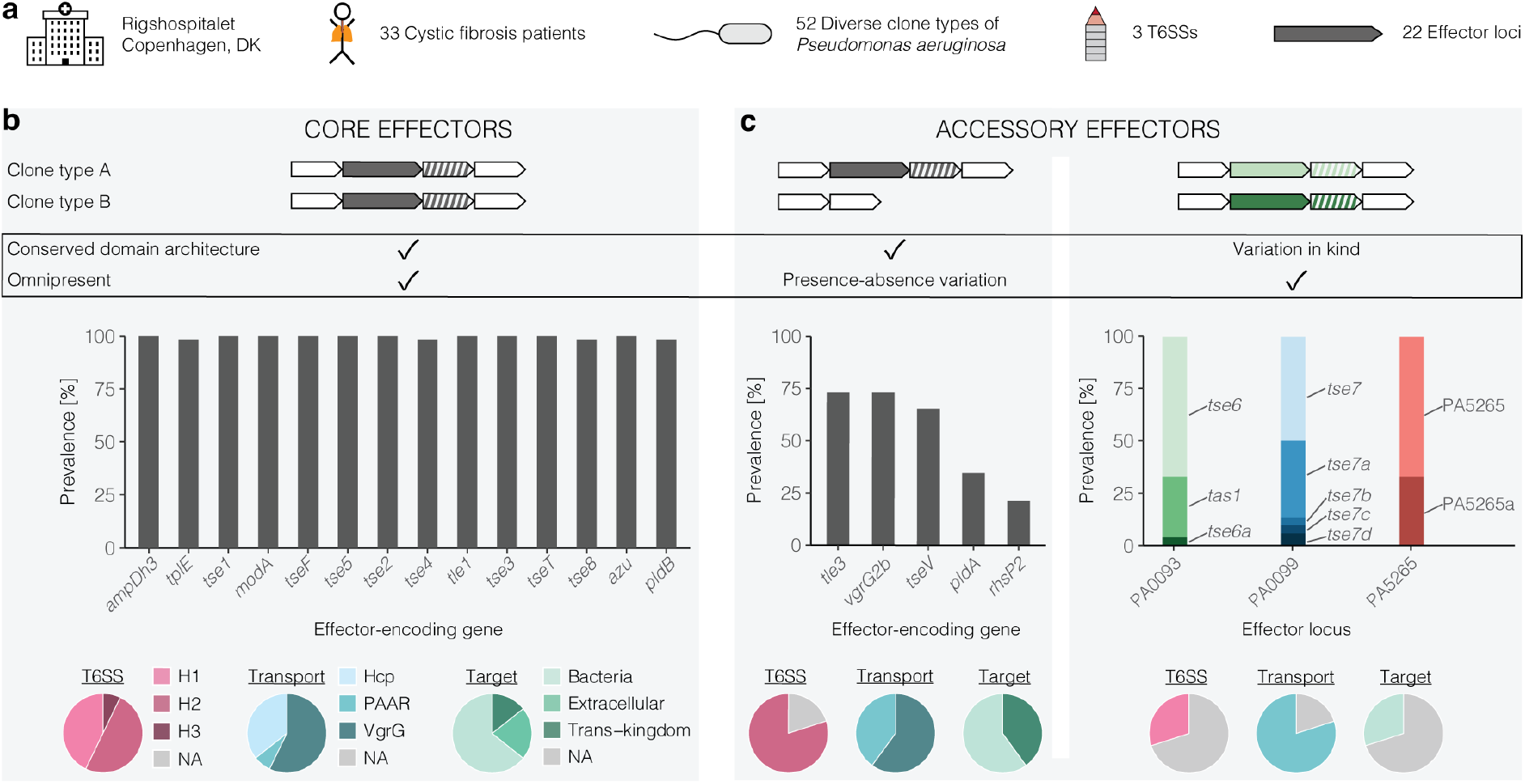
T6SS effectors are either omnipresent and conserved or show presence-absence variation and variation in kind. **a**, Overview of the analysed clone types and the known T6SS effector-encoding genes in *P. aeruginosa*. **b, c**, Core effectors are omnipresent and have a conserved domain architecture, accessory effectors show presence-absence variation or vary in their catalytic site. Graphical depictions of T6SS effector-encoding genes (filled in grey), immunity protein-encoding genes (striped in grey), and neighbouring genes (filled in white). Bar graphs show the prevalence of a given effector-encoding gene. For orientation, effector loci are labelled with PA numbers of the reference strain PAO1. Pie charts provide information on the type of T6SS an effector is linked to, the mechanism of effector translocation, and the effector targets. NA, not yet known. **b**, The blue horizontal line indicates a prevalence of 95%. **c**, Each shade of a given colour represents an effector with less than 30% amino acid identity at the catalytic site compared to another variant of the same locus.

## Results

### Core effectors are omnipresent and conserved across clone types whereas accessory effectors are not

We found fourteen effector-encoding genes were present in at least 98% of the analysed clone types (Fig. 1b, Supplementary Fig. 1). Effectors encoded at all 14 loci had a conserved domain architecture and shared 84 to 100% amino acid identity in a global alignment (Supplementary Fig. 2a). Only a small fraction of analysed sequences carried putative loss-of-function mutations that resulted in a shortened amino acid sequence (2 cases out of 724). Based on the high similarity and high prevalence of these effectors among the diverse clone types, we labelled them ‘core effectors’. Among them were effectors transported by all three T6SSs in association with either Hcp, PAAR or VgrG proteins, representing all T6SS-dependent mechanisms of secretion (Fig. 1b). Core effectors target prokaryotes, eukaryotic cells, and extracellular nutrients, suggesting a broad target spectrum.

Unlike core effectors, we found multiple effector-encoding genes in a fraction of the analysed clone types only, and we therefore referred to them as ‘accessory effectors’ (Fig. 1c). Among them, five showed presence-absence variation (Fig. 1c, Supplementary Fig. 2b). T6SS effector-encoding genes at three loci (PA0093, PA0099, PA5265) differed between clone types and are referred to as ‘variation in kind’ (Fig. 1c, Supplementary Fig. 2c-e). Characterised accessory effectors either belonged to the H1- or H2- and not H3-T6SS, and were associated with the secretion systems via either PAAR or VgrG and not Hcp proteins (Fig. 1c). Effectors secreted by H3-T6SS or associated with Hcp are unlikely omitted at random (probability P < 0.05), suggesting that accessory effectors are preferentially secreted by certain T6SSs via specific secretory mechanisms. The known targets of accessory effectors were prokaryotic and eukaryotic cells.

Altogether, we found (i) core effectors that are omnipresent and conserved in their domain architecture, and (ii) accessory effectors that vary in presence-absence or kind. The lack of diversity in some effectors implies T6SS-mediated phenotypes would be universally present across members of the species. The existence of accessory effectors opens the possibility of intraspecific diversity of effector combinations on the level of the individual isolate or strain.

### Effector set diversity across clone-types

Next, we characterised how core and accessory effectors were distributed across clone types. To do this, we identified all T6SS effectors present in each genome to define an effector set for each clone type (Fig. 2a). Assuming a random mix-and-match between all accessory effectors, as many as 960 unique effector sets were possible (Fig. 2b). Instead, we found 36 distinct effector sets among the 52 isolates (Fig. 2b, Supplementary Table 3). Each of these isolates belonged to a different clone type, capturing the phylogenetic diversity of a total of 473 clinical isolates in our collection^34^. Out of this collection, we analysed 421 additional isolates of the same clone types for other distinct effector sets, but did not find any more.

**Figure 2.**
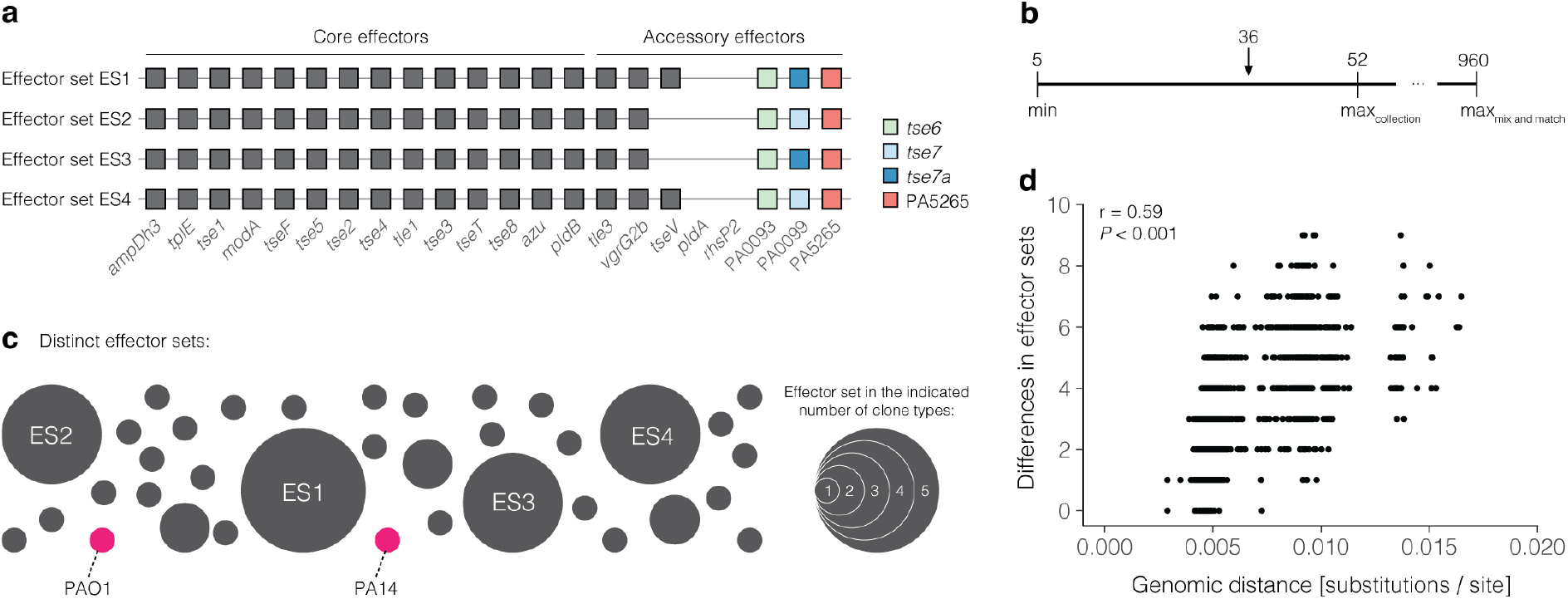
Fifty-two clone types harbour 36 distinct T6SS effector sets (ES). **a**, Schematic of four effector sets. Each set consists of all known effectors in a given genome. Each box represents one effector, shades of colour indicate different effector variants. **b**, Number of observed distinct effector sets (n=36) among the 52 clone types and a total of 960 theoretically possible distinct effector sets assuming mix and match. **c**, Total number of distinct effector sets among the analysed clone types. Each bubble represents one effector set, the size of the bubble depends on the number of clone types with a respective effector set. Bubbles of effector sets that are also found in lab reference strains are coloured in pink. Distribution of effector sets among clone types is not random (Monte Carlo simulation, *P* < 0.001). **d**, The correlation between the differences in effector sets (y-axis) and the genomic distance based on whole genomes (x-axis) was tested with a Pearson’s correlation coefficient.

Next, we analysed the relative abundance of the different effector sets, which are not randomly distributed across clone types (Monte Carlo simulation, *P* < 0.001). We found 29 effector sets in one clone type only and seven effector sets in at least two different clone types (Fig. 2c). The most abundant effector set (named ‘ES1’) was found in five clone types and consisted of the fourteen core effectors, and six accessory effectors (Fig. 2a, c). Of note, the four most abundant effector sets differed from each other only at two loci (presence-absence variation of *tseV* and variation in kind in PA0099) and were otherwise identical. Although some clone types had the same effector sets as the reference strains PAO1 and PA14, these sets were only found in one clone type each (highlighted in pink, Fig. 2c).

To test if the effector sets of clone types were indicative of their relatedness between each other, we compared the phylogenetic distances between clone types with their effector sets. We found a positive correlation (Pearson’s correlation coefficient 0.59, *P* < 0.001, 95% confidence interval 0.56 to 0.61) between differences in the T6SS effectors and differences in the whole genome (Fig. 2d). Two closely related clone types were significantly more likely to have similar effector sets than two distantly related clone types. This result might reflect the gradual diversification of the effector sets during the diversification of the species.

In summary, we demonstrated (i) that multiple diverse effector sets exist, (ii) which of the theoretically possible effector combinations are found in the isolate collection, and (iii) that some effector sets are more common than others. These findings indicate that the varying effectors might provide one clone type with an advantage over another clone type.

### Effector sets with the accessory effector PldA have a competitive advantage

To determine the minimal and maximal number of effectors per set, we analysed the total number of effectors in each of the 36 distinct sets. We found effector sets with as few as 17 and as many as 22 effectors (Fig. 3a). Of note, effector sets of the same size can differ in their effector composition, for example by accessory effectors that vary in kind. The frequencies of distinct effector sets were binomially distributed across the number of effectors per set *(P >* 0.05, Fig. 3a, Supplementary Fig. 3a), indicating that most sets had a size of 19 or 20 effectors whereas relatively few sets were small (17 to 18 effectors) or big (21 to 22 effectors). When taking into account that some effector sets are found in more than one clone type, effector sets with 19 and 20 effectors are also most common among the 52 distinct clone types (Fig. 3b). These results show that effector sets differ in size, and an intermediate number of effectors is most common among distinct effector sets and clone types.

**Figure 3.**
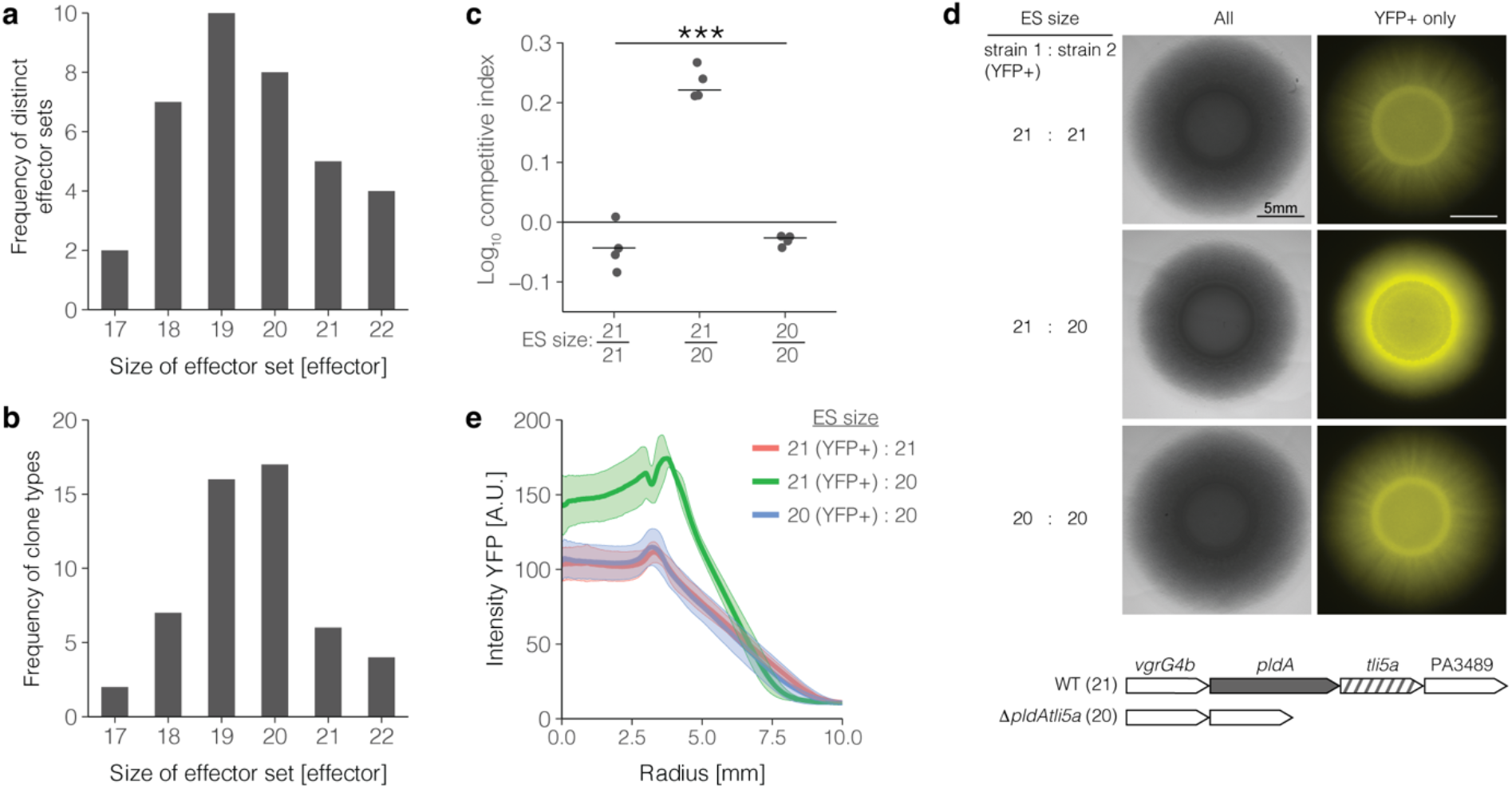
Effector set with additional anti-prokaryotic effector mediates bacteria-bacteria killing. **a**, Frequency distribution of effector sets of a certain size. **b**, Frequency distribution of clone types with a given size of effector sets. **c-e**, Competition experiments between two variants of the same strain with differing T6SS effector sets. Presence and absence of the accessory effector-encoding genes *pldA* results in an effector set of 20 and 21 effectors. Indicated strains were mixed at a 1:1 ratio and spotted onto agar plates. The experiment was performed four times. **c**, Strain with a bigger effector set outnumbers strain with a smaller effector set. Bars represent the mean ratio of bacterial counts (± SD) with ANOVA (\*\*\**P* < 0.001). **d**, Spatial distribution of the marked strain within the community of two competing strains (scale bars, 5mm). Representative images of one of four experiments are shown. **e**, Radial profiles (mean ± SD of four experiments) of the fluorescently marked strain starting at the centre towards the periphery of the community.

Effectors with presence-absence variation have previously been shown to be translocated with the T6SS and identified as toxins with anti-bacterial activity^45^. To test the difference between having one effector more or less in an effector set, we chose the accessory effector PldA that is present in some effector sets and absent from others. Competition experiments were performed between two variants of the same strain that only differ in this one effector of the otherwise identical effector set. Because effector protein-encoding genes are accompanied by immunity protein-encoding genes^1^, the presence or absence of an effector also reflects the presence or absence of its cognate immunity protein in our dataset (Supplementary Fig. 3b, c). A strain with a PldA-containing effector set of 21 effectors (here PAO1 wild-type) was observed to kill an otherwise identical strain with a PldA-deficient effector set of 20 effectors (here PAO1Δ*pldAtli5a*) (Fig. 3c). No killing was observed when two strains with the same effector sets were competed against each other (Fig 3c). This result demonstrates the advantage of a bigger, PldA-containing effector set in outnumbering a competing strain with a smaller, PldA-deficient effector set and confirms previous findings on the anti-prokaryotic activity of this ^45^ accessory effector.

T6SS-mediated killing is known to affect the spatial organisation of bacteria within communities^46^. To characterise the gain in space mediated by the PldA-containing effector set, we performed microscopic analysis of the two competing strains with a PldA-containing bigger effector set and a PldA-deficient smaller effector set (here PAO1 and PAO1Δ*pldAtli5a*). An expanding spot enables the observation of the spatial distribution of bacteria in an established model of a mixed community^46,47^. Therefore, mixed bacteria were spotted onto an agar surface and analysed for their spatial organization. We found that the strain with a PldA-containing, bigger effector set strongly dominated the centre and the periphery of the community (Fig. 3d). Radial profiles of fluorescence intensity across the community (Fig. 3e) further quantified the changes in spatial distribution and demonstrated the resulting advantage of the accessory effector PldA.

Taken together, we now know (i) the average size of effector sets in our dataset and (ii) that a larger effector set with the accessory effector PldA can mediate a competitive advantage in bacterial numbers and space.

### Differences in accessory effectors are detected between isolates of the same patient

Next, we tested whether intraspecific diversity in T6SS effectors, which we observed in a population of clinical isolates of an entire patient cohort from one geographic location, is also detected in a bacterial population of an individual patient. When analysing clone types that had been isolated from the same patient simultaneously, we find differences in their accessory effectors (Fig. 4). Clone types DK01, DK15, and DK53 from patient PID12139 differ in four of the accessory effectors with presence absence variation (such as *pldA*) and three accessory effectors that vary in kind. In patient PID61790, variation is observed in three accessory effectors with presence absence variation. In patient PID08136, four accessory effectors differ between clone types DK02 and DK20. These data show that T6SS effector sets not only differ between patients but also within patients. Whether the clone types were found in the same patient because or despite their different effector sets, which form the genetic basis also for T6SS-mediated killing between the clone types, remains unknown for now.

**Figure 4.**
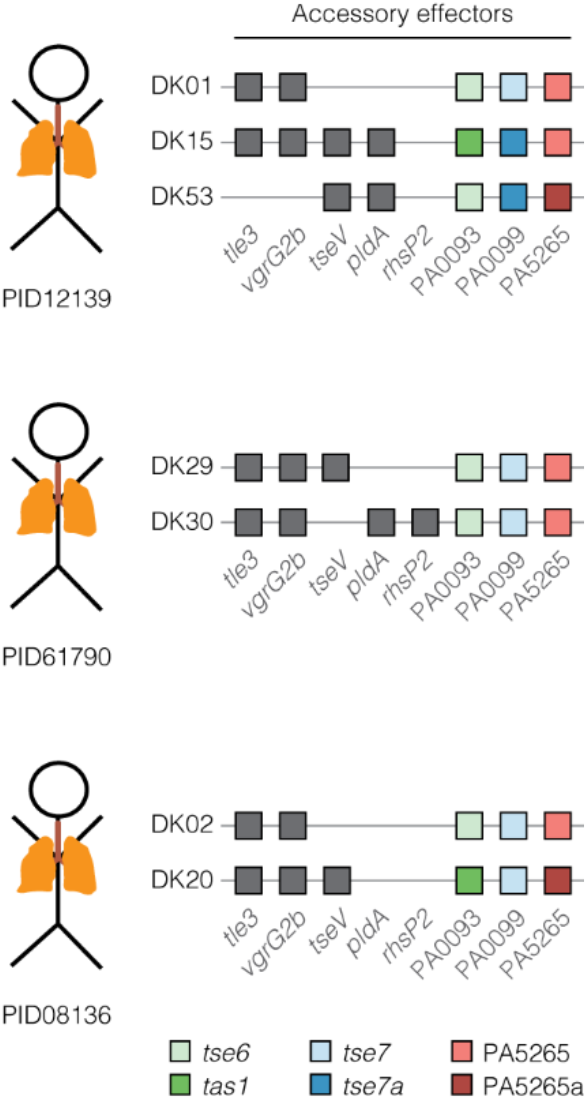
Simultaneously isolated clone types from the same patient differ in their effector sets. Schematic of the clone types’ accessory effectors and the patient they were isolated from. Each box represents one accessory effector. Grey boxes refer to effectors with presence absence variation and coloured boxes refer to effectors that vary in kind.

### Core effectors are omnipresent across isolate collections and contribute to the virulence of P. aeruginosa in vivo

Having established the diversity of T6SS effector sets, we decided to test whether our observations on core effectors were generalisable across isolates of various geographic regions and sources. Genome sequences of twenty diverse isolates collected from across the world from various clinical and environmental sources^48^ were analysed. We found all 14 core effectors in each of those isolates (Fig. 5a, Supplementary Fig. 4a). The genes were mostly intact and only very few sequences with putative loss-of-function mutations were detected that resulted in truncated amino acid sequences (2 cases out of 276 analysed sequences) (Supplementary Fig. 4a). Further, we analysed over 200 whole-genome sequences of *P. aeruginosa* available on NCBI, and found core effectors with a prevalence of at least 95% (Fig. 5a, Supplementary Fig. 4b). Two percent of the isolates lacked at least five core effectors; this may be attributed to the fact that they belonged to a distinct phylogenetic group (Supplementary Fig. 4c). Only few sequences contained putative loss-of-function mutations (40 cases out of 2937 analysed sequences). These findings (i) show a very high prevalence of core effectors across the species and (ii) suggest that they may be functional in the vast majority and confer a broad benefit.

**Figure 5.**
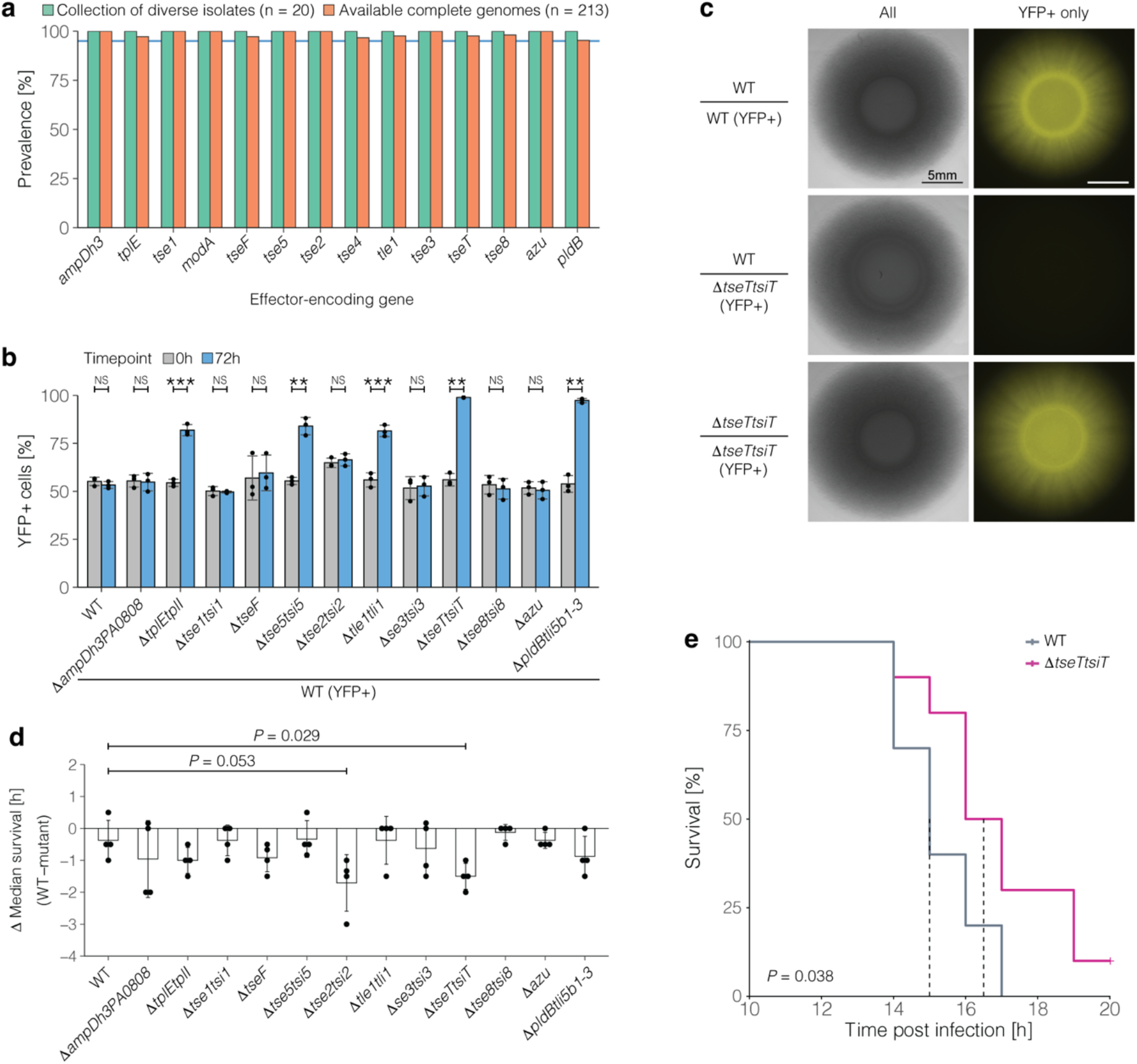
Core effectors mediate bacterial killing and contribute to the virulence of *P. aeruginosa*. **a**, Prevalence of core effectors in diverse isolate collections^48^. The blue horizontal line indicates a prevalence of 95%. **b**, Competition experiments to assess bacteria-bacteria killing between the indicated strains. Mean and standard deviation of three independent experiments are shown. Welch two-sample *t*-test was used to evaluate statistical significance (\*\*\**P* < 0.001; \*\**P* < 0.01; NS, not significant). **c**, Killing by TseT results in nearly complete eradication of sensitive strain. **d**, Virulence mediated by core effectors in *Galleria mellonella* infection experiments. Larvae were infected in the last left proleg with the indicated strain or PBS as a control. Mean and standard deviation of four independent experiments (each with 10 larvae) are shown. Each dot indicates the result of one experiment. Statistical significance was tested with a Welch two-sample *t*-test. **e**, Larvae infected with the core effector mutant Δ*tseTtsiT* live significantly longer than larvae infected with the wild type strain PAO1. Dotted lines indicate median survival times. Log-rank test was used for statistical evaluation.

Core effectors are known to be translocated with the T6SSs and most are known to mediate bacteria-bacteria killing^1,24–30,49–51^. Those observations were often made after the introduction of mutations to artificially activate the secretion systems *in vitro*, which is useful to study the T6SS molecular biology but might exaggerate phenotypes and is therefore of limited value to understand the ecological impact of core effectors. To test the role of each core effector for bacterial killing without artificially activating the secretion systems, we generated single-deletion mutants of twelve core effectors and, if present, their corresponding immunity protein-encoding genes (Supplementary Fig. 5). We find that the effector TseT (PA3907), with predicted endonuclease activity^24^, mediated the biggest competitive advantage among all core effectors under the conditions tested (Fig. 5b). Upon microscopic analysis of the community of competing bacteria, we observed nearly complete eradication of the strain sensitive to TseT-mediated killing (Fig. 5c). These results add value to the field by (i) demonstrating bacterial killing by core effectors without introducing genetic modifications to activate the secretion systems, (ii) providing a side-by-side comparison between effectors that have previously been studied independently by different laboratories in slightly different experimental set-ups and (iii) showing the power of core effectors in bacterial killing by reducing the competitor’s number and gaining space in a mixed community.

To test the contribution of core effectors to bacterial virulence *in vivo*, we used an established infection model of *Galleria mellonella*. We note that this infection model does not aim to reflect a specific human disease here but rather to improve our understanding of T6SS-mediated virulence in an organism more broadly. Although mutants with dysfunctional H1-T6SS and H3-T6SS have previously shown attenuated virulence upon systemic infection of *G. mellonella*^16,52^, the contribution of individual core effectors to the virulence of *P. aeruginosa* remains mostly unclear. We systemically infected *G. mellonella* larvae with mutants lacking the core effectors and recorded survival times. We observed less virulence for a mutant that lacks the putative endonuclease TseT, which had only been tested for anti-prokaryotic activity so far (Δ*tseTtsiT*; Fig. 5d, e; *t*-test *P* = 0. 029). We also observed on average a prolonged survival time for the mutant lacking *tse2* (Δ*tse2tsi2*; Fig. 5d; *t*-test *P* = 0.053), which encodes a putative mono-ADP-ribosyltransferase that has been reported as cytotoxic when expressed in mammalian cells *in vitro*^1,53^. These findings (i) present a first indication for the role of the core effector TseT during infection, (ii) show that core effectors contribute to the virulence of *P. aeruginosa*, and (iii) highlight the relevance of those effectors for bacterial pathogenesis.

## Discussion

Here, we report the diversity of T6SS effector sets in a population of clinical *P. aeruginosa* clone types and present evidence for the different roles of core and accessory effectors for the intraspecific diversity of this bacterial pathogen (Fig. 6). Our results show that caution should be taken when generalising conclusions about T6SS-mediated phenotypes based on the analysis of few strains: while we show on clinical isolates that combinations of accessory effectors differ considerably between clone types and enable intra-specific killing between *P. aeruginosa* bacteria, we found that core effectors with nutrient-acquiring, anti-prokaryotic, and anti-eukaryotic activity are indeed highly prevalent among strains even beyond our isolate collection. As such, *P. aeruginosa* stands out in comparison to other species that show intraspecific diversity in all known T6SS effectors^13,54–56^ or do not encode a T6SS in all strains^55,57,58^.

**Figure 6.**
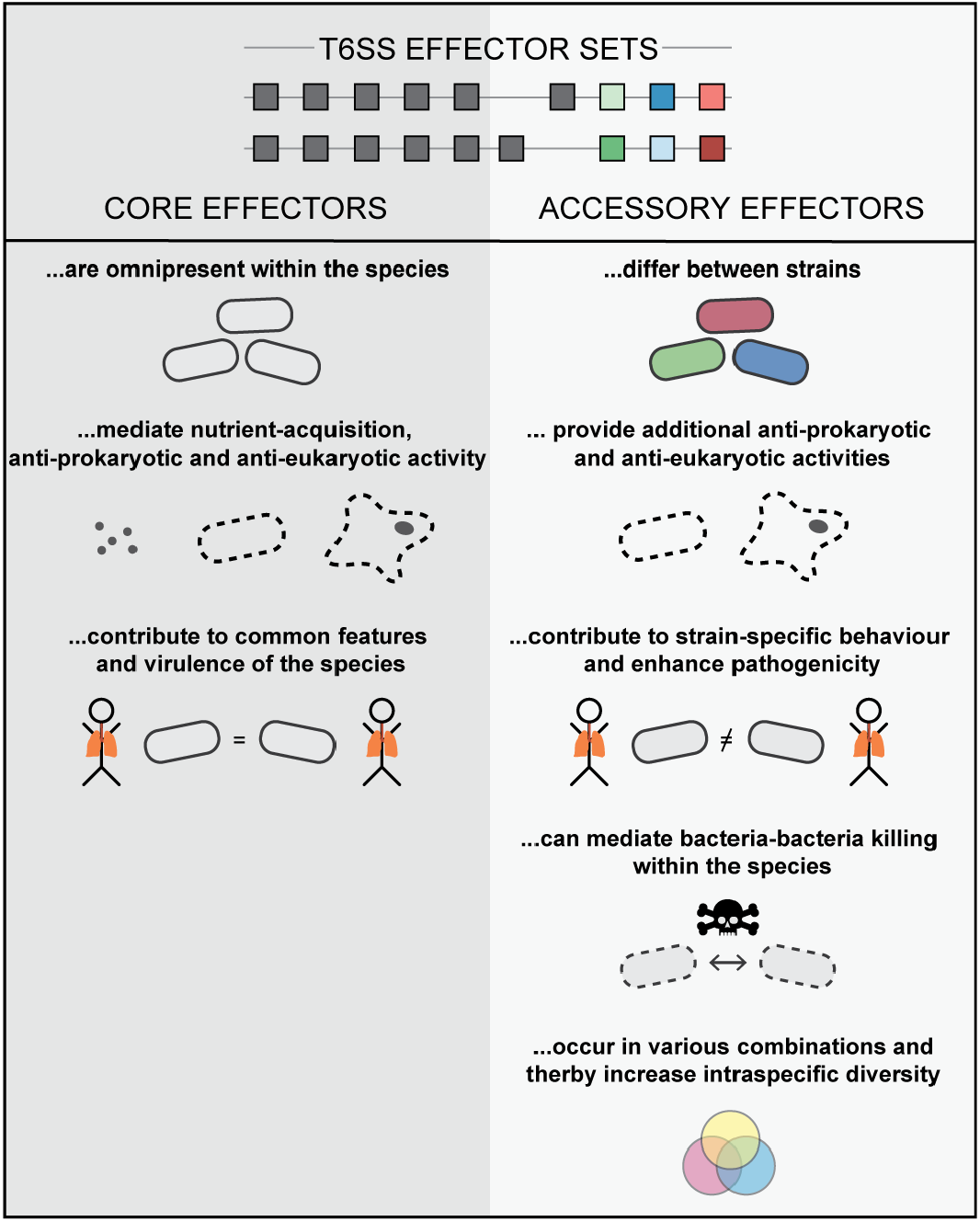
T6SS effector sets of *P. aeruginosa* are composed of core and accessory effectors. Core effectors (i) are highly prevalent within the species, (ii) mediate nutrient-acquisition, anti-prokaryotic, and anti-eukaryotic activity, and (iii) equally contribute to behaviour and virulence across strains of the species. In contrast, accessory effectors are less prevalent and differ between strains. They provide additional anti-prokaryotic and anti-eukaryotic activities. As a consequence of different accessory effectors, strains might differ in their behaviour and are more virulent. Two strains with differing accessory effectors can engage in T6SS-mediated killing of each other. The various combinations in which accessory effectors occur result in a multitude of diverse effector sets.

We propose that the herein described core effectors are among the few hundred genes in the core genome of *P. aeruginosa*, which has a pangenome of over 50,000 genes in total^59^. Many of these rare genes of the core genome are considered essential and fulfil housekeeping functions required for the growth of a bacterial cell^60^. Core effectors are not essential for bacterial survival in rich laboratory growth media, as was shown in here and by others^1,9,24,25,27–30,33,45,51,61,62^. Nevertheless, the high prevalence of the core effectors might be an indication for the importance of their function in the bacterium’s natural environment. Outside of the laboratory, *P. aeruginosa* faces diverse microbial competitors and scarce nutrient conditions, in which the anti-prokaryotic and nutrient-acquiring activities of core effectors might be universally beneficial to bacteria of this species. Although individual strains might differ in the regulation of core effectors, the genes are present across strains. We argue that our findings on virulence mediated by at least one core effector highlights the clinical relevance of the T6SSs for strains across the species.

Accessory effectors are some of many genes that differ between strains and contribute to strain-specific behaviour and pathogenicity. We found variation in accessory effectors between strains of the same and of different patients. Having or not having a certain accessory effector, like PldA, might affect the virulence of a particular strain. This notion is supported by experimental work showing PldA-mediated activation of the PI3K/Akt pathway of eukaryotic cells^14^ and by an association between an increased prevalence of *pldA* and a higher risk of exacerbations in non-CF bronchiectasis patients^21^. Further, strains with *pldA* were associated with acute pulmonary infections and multi-drug resistance^15^. However, we warn from measuring an isolate’s virulence by looking at T6SS effectors only. A PA7-related clinical isolate had previously been expected to be less virulent based on its lack of a type III secretion system but turned out more virulent because it had acquired another toxin^63^.

The herein described diversity of effector sets with various combinations of accessory effectors provides the genetic basis for extensive T6SS-mediated killing between *P. aeruginosa* strains. As observed in other species, strains with the same effector sets provide immunity to each other’s effectors and are considered compatible, whereas strains with different effector sets kill each other and cannot co-exist in a mixed community^54^. In this scenario, even seemingly redundant accessory effectors with a similar enzymatic activity to core effectors could confer a benefit. The accessory effector PldA and the core effector PldB are both lipases, which might be redundant when interacting with a bacterium outside the species^14^ and could be one reason for PldA not being present in all strains. However, the respective immunity proteins are specific to PldA or PldB, so that a *P. aeruginosa* strain with a PldA-containing effector set is able to kill a strain with an otherwise similar effector set that lacks PldA and the respective immunity protein^14^. Even if PldA is substituted by another effector that is yet unknown and encoded elsewhere in the genome, the two strains likely remain incompatible. Considering that additional T6SS effectors are still being discovered, the effector sets will likely become bigger and even more diverse in the upcoming years.

Our findings on diverse effector sets give hope for an applied use of the secretion system as a protein delivery tool. Recent attempts to engineer proteins for transport by the T6SS turned out challenging^26,64,65^. We showed that the effector sets that exist in our collection do not comprise effectors at random. Among the multiple mechanisms by which effectors are associated with the T6SS, we found Hcp-associated transport in the inner tube of the secretion system exclusively among core effectors. In contrast, the tip of the T6SS allows for transport of diverse accessory effectors with a PAAR domain, suggesting PAAR-mediated transport as the method of choice when developing T6SS delivery platforms^66^ and associating diverse engineered effectors with the secretion system.

## Materials and Methods

### Bacterial strains and growth conditions

*P. aeruginosa* was grown in LB broth at 37ºC. A list of strains used in this study is provided in Supplementary Table 4. In-frame deletions were generated via homologous recombination. In brief, fragments upstream and downstream of the gene of interest were PCR-amplified with primers p1, p2 and p3, p4 (all primers are listed in Supplementary Table 5). One continuous fragment was generated out of the two overlapping fragments in a second PCR and ligated into the vector pME3087^67^. The recombinant plasmid was transferred into the respective recipient strain during mating. Upon serial growth, tetracycline-sensitive cells were screened, mutants verified by PCR and confirmed by Sanger Sequencing. Fluorescently marked strains were generated as described by Schlechter *et al*.^68^ with the plasmid pMRE-Tn7-143 provided by addgene (#118495). Tetracycline was used at concentrations of 100µg ml^-1^.

### Competition in expanding colony

Bacterial strains were grown in liquid broth overnight, sub-cultured 1:100 on the day of the experiment and grown to early mid-logarithmic phase. Two strains were mixed at equal numbers to a final concentration of 5×10^7^ bacteria/30µl, of which 10µl were spotted onto LB agar plates (1.5% w/v). Plates were incubated at room temperature for 72h. During this time, plates were kept in a plastic box to prevent the agar from drying out.

### Microscopic imaging

An Axio Zoom.V16 microscope (Zeiss) with a PlanApo Z 0.5X objective was used to take brightfield and fluorescent images of the macrocolonies. Images were processed with Fiji^69^. Intensity profiles were created with the plug-in “Radial Profile Plot” and visualised using R.

### Flow cytometry

The BD Accuri C6 flow cytometer was used to quantify individual strains in the mixed community. Therefore, bacteria were scraped off the agar plate, resuspended in PBS and diluted. Threshold for the parameters forward scatter was set at 3,000 and for side scatter at 1,000. 10,000 events were quantified per sample. The competitive index was calculated by dividing the percentage of yellow fluorescent protein (YFP) cells per spot at 72h by the percentage of YFP cells at the start of the experiment.

### Growth curves

Overnight cultures were diluted 1:100 in LB, grown to exponential phase, further diluted and transferred into a 96-well plate to a starting OD600 of 0.001. The plate was incubated at 37 ºC, shaken at 162rpm, and OD600 measurements were taken every 15 minutes (Tecan, Spark). Three independent experiments were performed for each strain.

### *Galleria mellonella* infection assay

*G. mellonella* 6^*th*^- instar larvae were purchased from Faunatopics GmbH (Marbach, Germany). Upon arrival, larvae were stored at 10°C, for a maximum of 3 weeks. One day prior to infection, healthy looking, motile larvae with a body weight of 350 - 450 mg and without any signs of melanisation were selected and acclimatised at room temperature.

Infection assays of *G. mellonella* larvae were performed as previously described by McCarthy *et al*.^70^ with some modifications. In brief, *P. aeruginosa* overnight cultures were sub-cultured 1:50 in fresh LB medium and grown on a shaker at 37°C, 180 RPM until an *OD*_600_ of 0.6. Bacteria were pelleted by centrifugation, resuspended in sterile PBS, and serially diluted to 10^−4^. 10 µl bacterial suspension containing approximately 60 bacterial cells or 10 µl PBS alone (mock control) were injected into the last left proleg of larvae (n = 10 per treatment) using a Hamilton syringe. Larvae were stored at 4°C until all injections were completed and then transferred to a 37°C incubator. Twelve hours post infection, *G. mellonella* larvae were monitored on an hourly basis and checked for unresponsiveness and death. *P. aeruginosa* inocula were plated onto LB agar plates and enumerated for each experiment. Survival data is depicted as Kaplan-Meier curves. Kaplan-Meier curves of different conditions were compared using the log-rank test.

### Data accession

Genome assemblies for isolates belonging to the Copenhagen collection^34^ were downloaded from the European Nucleotide Archive (ENA). See Supplementary Table 1 for the accession codes of individual isolates. The accession codes for the reference strains PAO1 and PA14 are NC_002516 and CVON00000000, respectively. Raw reads of the 20 most common clones ^48^ were downloaded from the Sequence Read Archive (SRA). See Supplementary Table 6 for detailed accession information. All whole genomes of *P. aeruginosa* strains that were available on ENA by December 2020 were downloaded. Accession codes are found in Supplementary Table 7. Phylogenetic analysis of whole genomes was performed using andi^71^.

### *De novo* assembly

Raw reads were trimmed using BBDuk^72^ (Version 38.37) with default settings expect for the minimum quality, which was set to 20. Trimmed reads were *de novo* assembled into scaffolds using SPAdes^73^ (version 3.13.0) with default settings.

### T6SS effector-encoding gene analysis

Nucleotide sequences encoding for known *P. aeruginosa* T6SS effectors (Supplementary Table 2) (in case of *vgrG2b* only nucleotides 2271 to 3060 were used, which encode the enzymatically active domain) were extracted from the annotated reference strains PAO1 and PA14. These sequences were used as query sequences in a local blastn search against the contigs from the Copenhagen collection, assembled scaffolds from the most common clones, and all publicly available whole genomes to determine the prevalence of effector-encoding genes in the respective datasets. Nucleotide identities were calculated as described by Rohwer *et al*.^74^. The absence of genes was confirmed by analysing neighbouring genes. Additionally, sequences were manually inspected with Geneious^75^ (version 2019.2.3). Nucleotide sequences were translated, aligned to the PAO1 reference sequence using the Geneious^75^ alignment algorithm with default settings (version 2019.2.3), and the amino acid sequence identity was calculated as the percentage of residues that are identical to the reference. Effector variants share an amino acid sequence similarity of less than 30 % in the domain with the catalytic site^76–78^. The length of intact amino acid sequences was analysed as a read out for loss-of-function mutations by premature stop codons or frameshift mutations. Combinatorial analysis and stochastics were used to test if the distribution of core and accessory effectors is random among the associated T6SS, the mechanism of transport or their target.

## Acknowledgments

We thank Yasmin Claussen and Florian Schröder for technical assistance, Bernhard Haubold for discussions, and Rahul Unni, Sabrina Jabs, and Petra Bacher for comments on the manuscript. This work was funded by a European Molecular Biology Organization Long-Term Fellowship (ALTF 80-2015), the Daimler and Benz Foundation (grant 32-10/18), the German Cystic Fibrosis Association and the German Federal Ministry for Education and Research (grant 01KI2020) (to DU). AH receives support from the International Max Planck Research School for Evolutionary Biology.

## Author contributions

DU, ASG, MG, SBA, HKJ, SM, AH, VCV and AG contributed to the experimental design and provided advice, tools, and materials. AH and VCV performed the bioinformatics. OV generated bacterial mutants. AH, AG, and OV performed experiments. AH, AG, VCV and DU analysed the data. AH and DU wrote the first draft of the manuscript that was revised by all authors. DU and ASG conceived the project and supervised the study.

## References

1. Hood, R. D. et al. A type VI secretion system of Pseudomonas aeruginosa targets a toxin to bacteria. Cell Host Microbe 7, 25–37 (2010).

2. Pukatzki, S., Ma, A. T., Revel, A. T., Sturtevant, D. & Mekalanos, J. J. Type VI secretion system translocates a phage tail spike-like protein into target cells where it cross-links actin. Proc. Natl. Acad. Sci. U. S. A. 104, 15508–13 (2007).

3. Si, M. et al. Manganese scavenging and oxidative stress response mediated by type VI secretion system in Burkholderia thailandensis. Proc. Natl. Acad. Sci. 114, E2233–E2242 (2017).

4. Mougous, J. D. et al. A virulence locus of Pseudomonas aeruginosa encodes a protein secretion apparatus. Science (80-.). 312, 1526–1530 (2006).

5. Russell, A. B. et al. A type VI secretion-related pathway in Bacteroidetes mediates interbacterial antagonism. Cell Host Microbe 16, 227–236 (2014).

6. Speare, L. et al. Bacterial symbionts use a type VI secretion system to eliminate competitors in their natural host. Proc. Natl. Acad. Sci. 115, E8528–E8537 (2018).

7. Pukatzki, S. et al. Identification of a conserved bacterial protein secretion system in Vibrio cholerae using the Dictyostelium host model system. Proc. Natl. Acad. Sci. U. S. A. 103, 1528–33 (2006).

8. Russell, A. B., Peterson, S. B. & Mougous, J. D. Type VI secretion system effectors: poisons with a purpose. Nat. Rev. Microbiol. 12, 137–48 (2014).

9. Lin, J. et al. A Pseudomonas T6SS effector recruits PQS-containing outer membrane vesicles for iron acquisition. Nat. Commun. 8, 1–12 (2017).

10. Trunk, K. et al. The type VI secretion system deploys antifungal effectors against microbial competitors. Nat. Microbiol. 3, 920–931 (2018).

11. Le, N.-H., Pinedo, V., Lopez, J., Cava, F. & Feldman, M. F. Killing of Gram-negative and Gram-positive bacteria by a bifunctional cell wall-targeting T6SS effector. Proc. Natl. Acad. Sci. 118, e2106555118 (2021).

12. Russell, A. B. et al. A widespread bacterial type VI secretion effector superfamily identified using a heuristic approach. Cell Host Microbe 11, 538–549 (2012).

13. Kirchberger, P. C., Unterweger, D., Provenzano, D., Pukatzki, S. & Boucher, Y. Sequential displacement of type VI secretion system effector genes leads to evolution of diverse immunity gene arrays in Vibrio cholerae. Sci. Rep. 7, 1–12 (2017).

14. Jiang, F., Waterfield, N. R., Yang, J., Yang, G. & Jin, Q. A Pseudomonas aeruginosa type VI secretion phospholipase D effector targets both prokaryotic and eukaryotic cells. Cell Host Microbe 15, 600–610 (2014).

15. Boulant, T. et al. Higher prevalence of PldA, a Pseudomonas aeruginosa trans-kingdom H2-type VI secretion system effector, in clinical isolates responsible for acute infections and in multidrug resistant strains. Front. Microbiol. 9, 1–7 (2018).

16. Pissaridou, P. et al. The Pseudomonas aeruginosa T6SS-VgrG1b spike is topped by a PAAR protein eliciting DNA damage to bacterial competitors. Proc. Natl. Acad. Sci. U. S. A. 115, 12519–12524 (2018).

17. Ahmad, S. et al. An interbacterial toxin inhibits target cell growth by synthesizing (p)ppApp. Nature 575, 674–678 (2019).

18. LaCourse, K. D. et al. Conditional toxicity and synergy drive diversity among antibacterial effectors. Nat. Microbiol. 3, 440–446 (2018).

19. Gibson, R. L., Burns, J. L. & Ramsey, B. W. Pathophysiology and management of pulmonary infections in cystic fibrosis. Am. J. Respir. Crit. Care Med. 168, 918–951 (2003).

20. Perault, A. I. et al. Host adaptation predisposes Pseudomonas aeruginosa to type VI secretion system-mediated predation by the Burkholderia cepacia Complex. Cell Host Microbe 28, 534–547.e3 (2020).

21. Luo, R.-G. et al. Presence of pldA and exoU in mucoid Pseudomonas aeruginosa is associated with high risk of exacerbations in non–cystic fibrosis bronchiectasis patients. Clin. Microbiol. Infect. 25, 601–606 (2019).

22. Sana, T. G., Berni, B. & Bleves, S. The T6SSs of Pseudomonas aeruginosa strain PAO1 and their effectors: Beyond bacterial-cell targeting. Front. Cell. Infect. Microbiol. 6, 61 (2016).

23. Lesic, B., Starkey, M., He, J., Hazan, R. & Rahme, L. G. Quorum sensing differentially regulates Pseudomonas aeruginosa type VI secretion locus I and homologous loci II and III, which are required for pathogenesis. Microbiology 155, 2845–2855 (2009).

24. Burkinshaw, B. J. et al. A type VI secretion system effector delivery mechanism dependent on PAAR and a chaperone-co-chaperone complex. Nat. Microbiol. 3, 632–640 (2018).

25. Han, Y. et al. A Pseudomonas aeruginosa type VI secretion system regulated by CueR facilitates copper acquisition. PLoS Pathog. 15, e1008198 (2019).

26. Wettstadt, S., Wood, T. E., Fecht, S. & Filloux, A. Delivery of the Pseudomonas aeruginosa phospholipase effectors PldA and PldB in a VgrG-And H2-T6SS-dependent manner. Front. Microbiol. 10, 1–18 (2019).

27. Wang, T. et al. A type VI secretion system delivers a cell wall amidase to target bacterial competitors. Mol. Microbiol. 114, 308–321 (2020).

28. Nolan, L. M. et al. Identification of Tse8 as a type VI secretion system toxin from Pseudomonas aeruginosa that targets the bacterial transamidosome to inhibit protein synthesis in prey cells. Nat. Microbiol. 6, 1199–1210 (2021).

29. Hachani, A., Allsopp, L. P., Oduko, Y. & Filloux, A. The VgrG proteins are ‘à la carte’ delivery systems for bacterial type VI effectors. J. Biol. Chem. 289, 17872–17884 (2014).

30. Whitney, J. C. et al. Genetically distinct pathways guide effector export through the type VI secretion system. Mol. Microbiol. 92, 529–542 (2014).

31. Cianfanelli, F. R. et al. VgrG and PAAR proteins define distinct versions of a functional type VI secretion system. PLoS Pathog. 12, 1–27 (2016).

32. Howard, S. A. et al. The breadth and molecular basis of Hcp-driven type VI secretion system effector delivery. MBio (2021). doi:10.1128/mBio.00262-21

33. Wang, T. et al. Pseudomonas aeruginosa T6SS-mediated molybdate transport contributes to bacterial competition during anaerobiosis. Cell Rep. 35, 108957 (2021).

34. Marvig, R. L., Sommer, L. M., Molin, S. & Johansen, H. K. Convergent evolution and adaptation of Pseudomonas aeruginosa within patients with cystic fibrosis. Nat. Genet. 47, 57–64 (2015).

35. Andersen, S. B., Marvig, R. L., Molin, S., Krogh Johansen, H. & Griffin, A. S. Long-term social dynamics drive loss of function in pathogenic bacteria. Proc. Natl. Acad. Sci. 112, 10756–10761 (2015).

36. La Rosa, R., Rossi, E., Feist, A. M., Johansen, H. K. & Molin, S. Compensatory evolution of Pseudomonas aeruginosa’s slow growth phenotype suggests mechanisms of adaptation in cystic fibrosis. Nat. Commun. 12, 3186 (2021).

37. Andersen, S. B. et al. Diversity, prevalence, and longitudinal occurrence of type II toxin-antitoxin systems of Pseudomonas aeruginosa infecting cystic fibrosis lungs. Front. Microbiol. 8, 1–12 (2017).

38. Ghoul, M. et al. Bacteriocin-mediated competition in cystic fibrosis lung infections. Proc. R. Soc. B Biol. Sci. 282, 20150972 (2015).

39. Andersen, S. B. et al. Privatisation rescues function following loss of cooperation. Elife 7, (2018).

40. Rossi, E., Falcone, M., Molin, S. & Johansen, H. K. High-resolution in situ transcriptomics of Pseudomonas aeruginosa unveils genotype independent patho-phenotypes in cystic fibrosis lungs. Nat. Commun. 9, 3459 (2018).

41. Bartell, J. A. et al. Evolutionary highways to persistent bacterial infection. Nat. Commun. 10, (2019).

42. Gabrielaite, M., Johansen, H. K., Molin, S., Nielsen, F. C. & Marvig, R. L. Gene loss and acquisition in lineages of Pseudomonas aeruginosa evolving in cystic fibrosis patient airways. MBio 11, 1–16 (2020).

43. Bartell, J. A. et al. Bacterial persisters in long-term infection: Emergence and fitness in a complex host environment. PLOS Pathog. 16, e1009112 (2020).

44. La Rosa, R., Johansen, H. K. & Molin, S. Convergent metabolic specialization through distinct evolutionary paths in Pseudomonas aeruginosa. MBio 9, (2018).

45. Russell, A. B. et al. Diverse type VI secretion phospholipases are functionally plastic antibacterial effectors. Nature 496, 508–512 (2013).

46. McNally, L. et al. Killing by Type VI secretion drives genetic phase separation and correlates with increased cooperation. Nat. Commun. 8, 14371 (2017).

47. Kalziqi, A. et al. Immotile active matter: Activity from death and reproduction. Phys. Rev. Lett. 120, 018101 (2018).

48. Hilker, R. et al. Interclonal gradient of virulence in the Pseudomonas aeruginosa pangenome from disease and environment. Environ. Microbiol. 17, 29–46 (2015).

49. Hachani, A. et al. Type VI secretion system in Pseudomonas aeruginosa: Secretion and multimerization of VgrG proteins. J. Biol. Chem. 286, 12317–12327 (2011).

50. Russell, A. B. et al. Type VI secretion delivers bacteriolytic effectors to target cells. Nature 475, 343–347 (2011).

51. Silverman, J. M. et al. Haemolysin coregulated protein is an exported receptor and chaperone of type VI secretion substrates. Mol. Cell 51, 584–593 (2013).

52. Li, Y., Chen, L., Zhang, P., Bhagirath, A. Y. & Duan, K. ClpV3 of the H3-type VI secretion system (H3-T6SS) affects multiple virulence factors in Pseudomonas aeruginosa. Front. Microbiol. 11, (2020).

53. Robb, C. S., Robb, M., Nano, F. E. & Boraston, A. B. The structure of the toxin and type six secretion system substrate Tse2 in complex with its immunity protein. Structure 24, 277–284 (2016).

54. Unterweger, D. et al. The Vibrio cholerae type VI secretion system employs diverse effector modules for intraspecific competition. Nat. Commun. 5, 1–9 (2014).

55. Steele, M. I., Kwong, W. K., Whiteley, M. & Moran, N. A. Diversification of type VI secretion system toxins reveals ancient antagonism among bee gut microbes. MBio 8, (2017).

56. Santoriello, F. J., Kirchberger, P. C., Boucher, Y. & Pukatzki, S. Pandemic Vibrio cholerae acquired competitive traits from an environmental Vibrio species. bioRxiv (2021). doi:10.1101/2021.05.28.446156

57. Fridman, C. M., Keppel, K., Gerlic, M., Bosis, E. & Salomon, D. A comparative genomics methodology reveals a widespread family of membrane-disrupting T6SS effectors. Nat. Commun. 11, (2020).

58. García-Bayona, L., Coyne, M. J. & Comstock, L. E. Mobile Type VI secretion system loci of the gut Bacteroidales display extensive intra-ecosystem transfer, multi-species spread and geographical clustering. PLOS Genet. 17, e1009541 (2021).

59. Freschi, L. et al. The Pseudomonas aeruginosa pan-genome provides new insights on its population structure, horizontal gene transfer, and pathogenicity. Genome Biol. Evol. 11, 109–120 (2019).

60. Turner, K. H., Wessel, A. K., Palmer, G. C., Murray, J. L. & Whiteley, M. Essential genome of Pseudomonas aeruginosa in cystic fibrosis sputum. Proc. Natl. Acad. Sci. 112, 4110–4115 (2015).

61. Jiang, F. et al. The Pseudomonas aeruginosa type VI secretion PGAP1-like effector induces host autophagy by activating endoplasmic reticulum stress. Cell Rep. 16, 1502–1509 (2016).

62. Barret, M., Egan, F., Fargier, E., Morrissey, J. P. & O’Gara, F. Genomic analysis of the type VI secretion systems in Pseudomonas spp.: Novel clusters and putative effectors uncovered. Microbiology 157, 1726–1739 (2011).

63. Elsen, S. et al. A type III secretion negative clinical strain of Pseudomonas aeruginosa employs a two-partner secreted exolysin to induce hemorrhagic pneumonia. Cell Host Microbe 15, 164–176 (2014).

64. Wettstadt, S. & Filloux, A. Manipulating the type VI secretion system spike to shuttle passenger proteins. PLoS One 15, 1–20 (2020).

65. Wettstadt, S., Lai, E.-M. & Filloux, A. Solving the puzzle: Connecting a heterologous Agrobacterium tumefaciens T6SS effector to a Pseudomonas aeruginosa spike complex. Front. Cell. Infect. Microbiol. 10, (2020).

66. Jana, B., Keppel, K. & Salomon, D. Engineering a customizable antibacterial T6SS-based platform in Vibrio natriegens. EMBO Rep. (2021). doi:10.15252/embr.202153681

## Methods references

67. Scott, T. A., Heine, D., Qin, Z. & Wilkinson, B. An L-threonine transaldolase is required for L-threo-β-hydroxy-α-amino acid assembly during obafluorin biosynthesis. Nat. Commun. 8, 15935 (2017).

68. Schlechter, R. O. et al. Chromatic bacteria – A broad host-range plasmid and chromosomal insertion toolbox for fluorescent protein expression in bacteria. Front. Microbiol. 9, (2018).

69. Schindelin, J. et al. Fiji: an open-source platform for biological-image analysis. Nat. Methods 9, 676–682 (2012).

70. McCarthy, R. R. c-di-GMP Signaling. 1657, (Springer New York, 2017).

71. Haubold, B., Klötzl, F. & Pfaffelhuber, P. Andi: Fast and accurate estimation of evolutionary distances between closely related genomes. Bioinformatics 31, 1169–1175 (2015).

72. Bushnell, B. BBMap (BBDuk plugin). (2015).

73. Nurk, S. et al. Assembling single-cell genomes and mini-metagenomes from chimeric MDA products. J. Comput. Biol. 20, 714–737 (2013).

74. Rohwer, R. R., Hamilton, J. J., Newton, R. J. & McMahon, K. D. TaxAss: Leveraging a custom freshwater database achieves fine-scale taxonomic resolution. mSphere 3, 1–14 (2018).

75. Kearse, M. et al. Geneious Basic: An integrated and extendable desktop software platform for the organization and analysis of sequence data. Bioinformatics 28, 1647–1649 (2012).

76. Rost, B. Twilight zone of protein sequence alignments. Protein Eng. Des. Sel. 12, 85–94 (1999).

77. Orengo, C. A. & Thornton, J. M. Protein families and their evolution - a structural perspective. Annu. Rev. Biochem. 74, 867–900 (2005).

78. Yu, L. et al. Grammar of protein domain architectures. Proc. Natl. Acad. Sci. 116, 3636–3645 (2019).

