## Supplementary information for "Core and accessory effectors of type VI secretion systems contribute differently to the intraspecific diversity of *Pseudomonas aeruginosa*"

Supplementary File

This file includes:

Supplementary Figures 1-5.

Supplementary Tables 1, 2, 4, 5, 6, 7.

Supplementary Table 3 is provided as a separate .xls file.

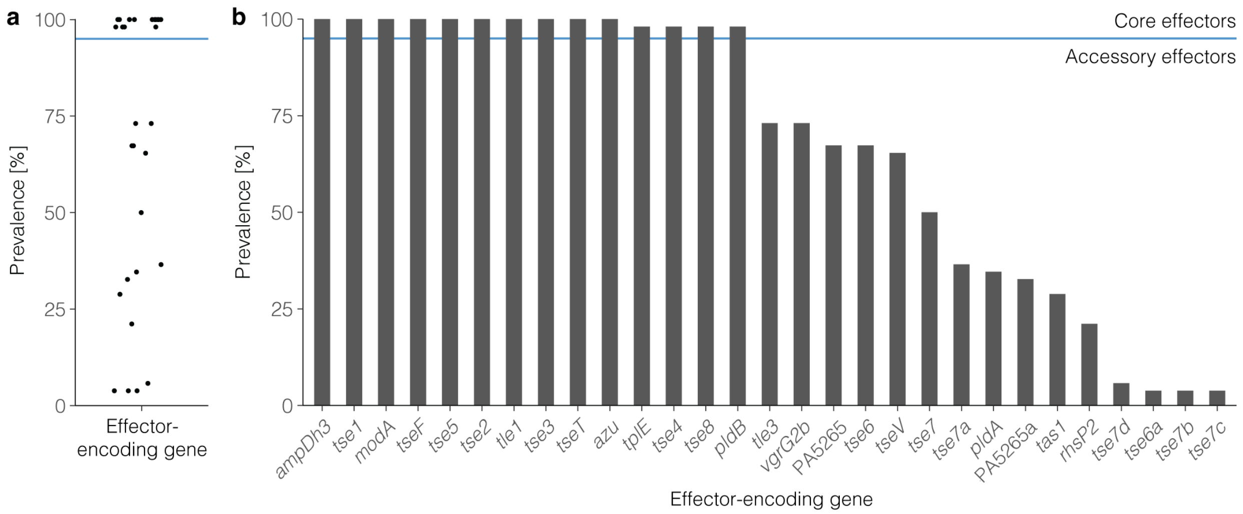


**Supplementary Figure 1.** Prevalence of T6SS effector-encoding genes among 52 clinical isolates of the Copenhagen collection. **a,** A group of genes with very high prevalence stands out from other genes with a lower prevalence. Each dot represents one effector-encoding gene. **b**, The same data than in panel A is shown as bar graphs to visualize the prevalence of individual effectors more easily. Effector-encoding genes are ordered by their prevalence. The blue horizontal line indicates a prevalence of 95 %.

**a**

| Effector locus | Minimal nucleotide identity | Minimal amino acid identity |
|---|---|---|
| PA0807 | 98.96 % | 99.61 % |
| PA1510 | 91.08 % | 91.92 % |
| PA1844 | 99.36 % | 99.35 % |
| PA1863 | 98.54 % | 98.41 % |
| PA2374 | 97.39 % | 96.63 % |
| PA2684 | 98.66 % | 84.50 % |
| PA2702 | 98.32 % | 98.73 % * |
| PA2774 | 98.64 % | 98.48 % * |
| PA3290 | 74.78 % | 90.35 % |
| PA3484 | 98.78 % | 99.26 % |
| PA3907 | 97.1 % | 99.24 % |
| PA4163 | 99.36 % | 99.42 % |
| PA4922 | 99.33 % | 99.32 % |
| PA5089 | 92.72 % | 92.08 % |

**b**

| Effector locus | Minimal nucleotide identity | Minimal amino acid identity |
|---|---|---|
| PA0260 | 84.94 % | 95.08 % |
| PA0262 | 63.8 % | 95.75 % |
| PA0822 | 75.33 % | 91.89 % |
| PA3487 | 94.37 % | 93.27 % |
| PA14_43100 | 99.44 % | 98.88 % * |

**c** Amino acid alignment: Tse6 (PA0093) variants

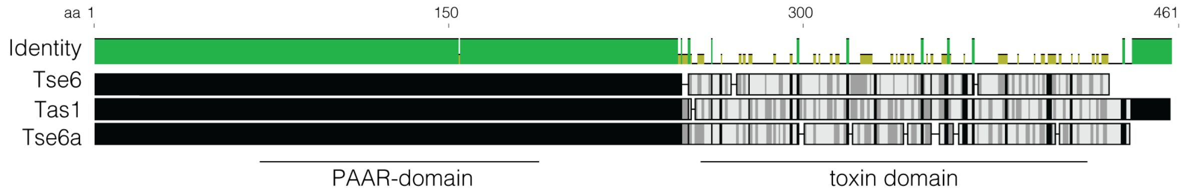


| | Tse6-tox | Tas1-tox | Tse6a-tox |
|---|---|---|---|
| Tse6-tox | | 9.38% | 11.88% |
| Tas1-tox | 9.38% | | 15.63% |
| Tse6a-tox | 11.88% | 15.63% | |

**d** Amino acid alignment: Tse7 (PA0099) variants

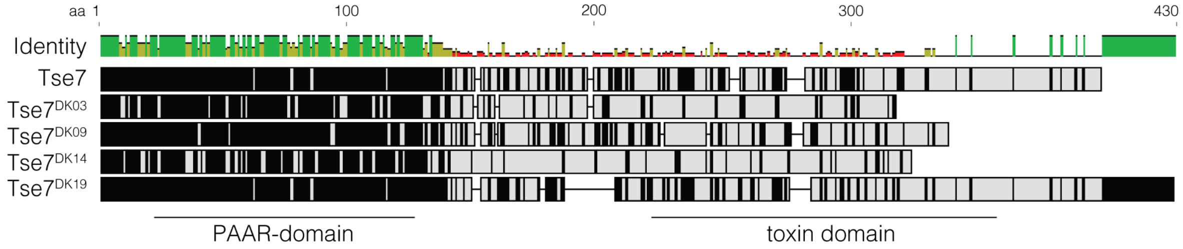


| | Tse7-tox | Tse7a-tox | Tse7b-tox | Tse7c-tox | Tse7d-tox |
|---|---|---|---|---|---|
| Tse7-tox | | 10.00% | 12.15% | 8.33% | 24.39% |
| Tse7a-tox | 10.00% | | 5.56% | 4.44% | 11.11% |
| Tse7b-tox | 12.15% | 5.56% | | 11.46% | 9.35% |
| Tse7c-tox | 8.33% | 4.44% | 11.46% | | 7.29% |
| Tse7d-tox | 24.39% | 11.11% | 19.35% | 7.29% | |

**e** Amino acid alignment: PA5265 variants

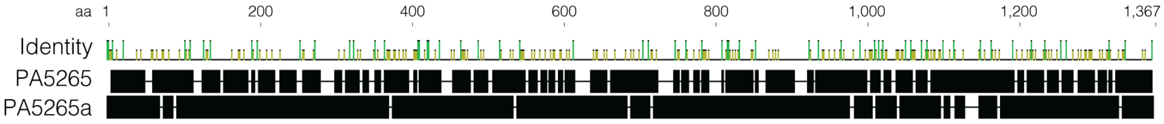


Amino acid identity: 18.0%

**Supplementary Figure 2.** Minimal amino acid identity resulting from comparisons of protein sequences of either core effector (**a**) or effectors with presence-absence variation (**b**) to the respective amino acid sequence of the reference strain PAO1. An asterisk indicates that one clone type was excluded from the amino acid alignment because of frameshift mutations in the effector-encoding gene. **c-e**, Amino acid alignments of effectors varying in kind and distance matrices indicating amino acid sequence similarities between the toxin domains of Tse6 (**c**), Tse7 (**d**), and PA5265 (**e**) variants.

a

| Isolate selection | p-value |
|---|---|
| Distinct effector sets | 0.06 |
| Clone types | 0.46 |

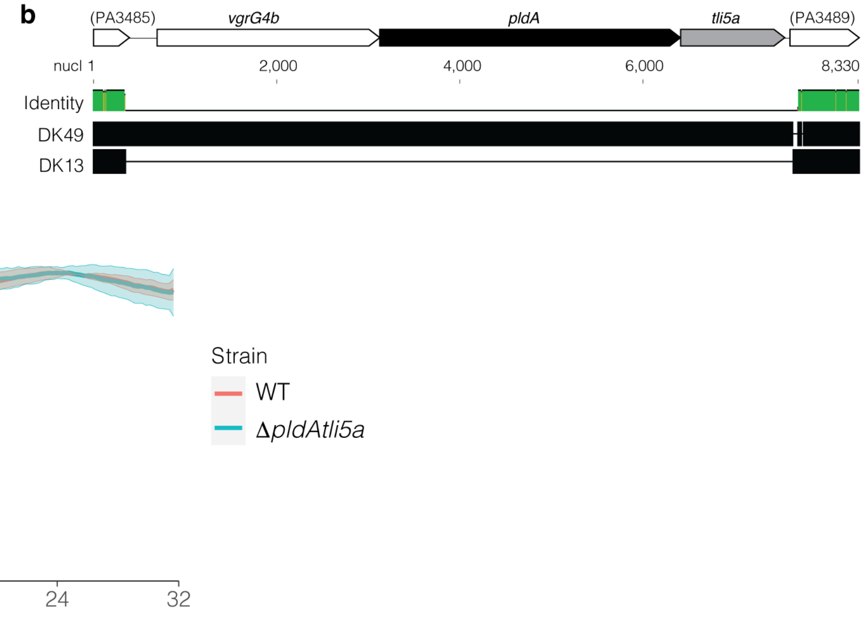


**Supplementary Figure 3**. **a**, P-values for $\chi^2$-test to test for a binomial distribution. **b**, Nucleotide alignment of the genomic region including *pldA* (PA3487) and *tli5a* (PA3488) of the two clone types DK49 and DK13. **c**, Growth curve of the strain with 21 effectors (PAO1 WT) and the strain with 20 effectors (PAO1 *ΔpldAtli5a)*. Data is presented as the mean ± SD of three experiments.

**a** Selection of diverse isolates (n = 20)

| Effector locus | Minimal nucleotide identity |
|---|---|
| PA0807 | 98.96 % |
| PA1510 | 91.08 % |
| PA1844 | 99.36 % |
| PA1863 | 98.54 % |
| PA2374 | 97.03 % |
| PA2684 | 89.38 % |
| PA2702 | 97.69 % |
| PA2774 | 96.77 % * |
| PA3290 | 74.82 % |
| PA3484 | 97.88 % |
| PA3907 | 96.31 % |
| PA4163 | 98.66 % |
| PA4922 | 98.66 % |
| PA5089 | 92.63 % |

**b** Available complete genomes (n = 213)

| Effector locus | Minimal nucleotide identity |
|---|---|
| PA0807 | 95.44 % |
| PA1510 | 65.6 % * |
| PA1844 | 92.99 % * |
| PA1863 | 94.18 % |
| PA2374 | 97.39 % |
| PA2684 | 88.14 % * |
| PA2702 | 97.48 % * |
| PA2774 | 98.64 % * |
| PA3290 | 73.47 % * |
| PA3484 | 96.74 % |
| PA3907 | 98.98 % * |
| PA4163 | 96.49 % * |
| PA4922 | 96.87% |
| PA5089 | 84.6 % * |

**c**

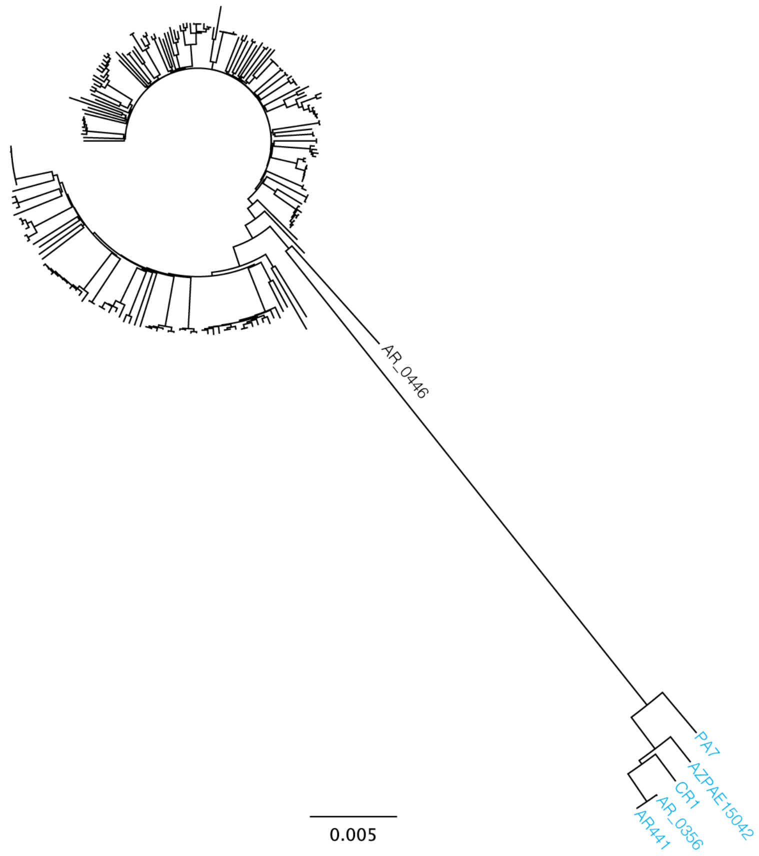


**Supplementary Figure 4.** Minimal nucleotide identity resulting from pairwise comparisons of core effector-encoding genes of diverse isolates (**a**) and of available complete genomes (**b**) to the respective gene of the reference strain PAO1. An asterisk indicates that at least one sequence was found with a mutation resulting in a premature stop codon. **c**, Phylogeny of the analysed whole genome sequences. Strains missing at least five of the same core effectors are indicated in blue.

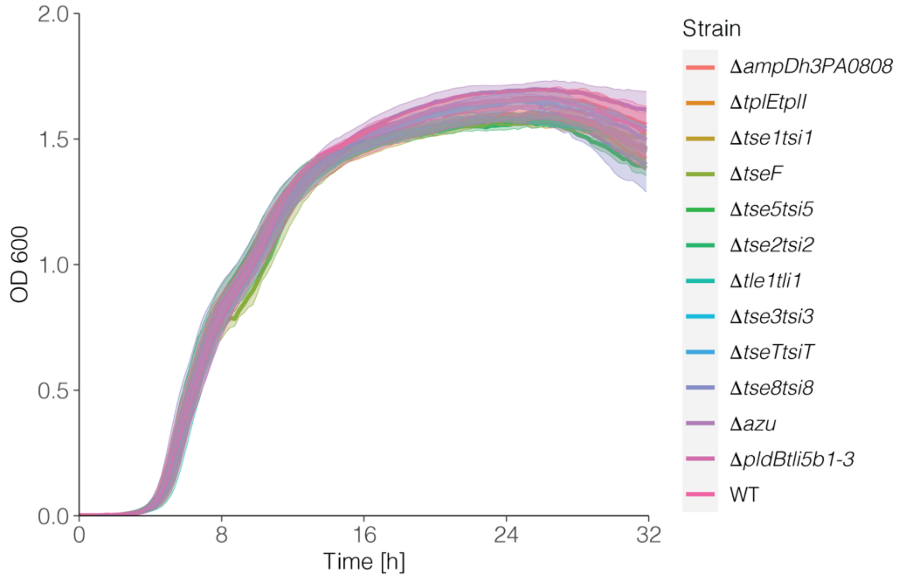


**Supplementary Figure 5.** Growth curves of PAO1 WT and all mutants deficient in a core effector-encoding gene (and if present, also the corresponding immunity protein-encoding gene). Data is presented as the mean ± SD of three experiments.

**Supplementary Table 1.** Isolates used for the genome analysis.

| Clone type | Isolate name | Patient origin | Sampling date | Sequence Read Archive (SRA) accession number |
|---|---|---|---|---|
| DK01 | 433 | PID12139 | 5/1/06 | ERS402771 |
| DK02 | 165 | PID08136 | 3/18/11 | ERS402479 |
| DK03 | 317 | PID31159 | 7/2/07 | ERS402641 |
| DK04 | 23 | PID09574 | 9/5/07 | ERS402554 |
| DK05 | 27 | PID09574 | 5/2/12 | ERS402588 |
| DK06 | 390 | PID42824 | 3/15/06 | ERS402720 |
| DK07 | 19 | PID51036 | 11/21/11 | ERS402507 |
| DK08 | 66 | PID23844 | 9/15/10 | ERS402795 |
| DK09 | 77 | PID20336 | 5/8/06 | ERS402800 |
| DK10 | 94 | PID21992 | 12/6/11 | ERS402819 |
| DK11 | 92 | PID21992 | 5/14/12 | ERS402817 |
| DK12 | 95 | PID76609 | 4/4/06 | ERS402820 |
| DK13 | 122 | PID83337 | 11/10/05 | ERS402421 |
| DK14 | 132 | PID35193 | 5/19/08 | ERS402436 |
| DK15 | 427 | PID12139 | 8/29/06 | ERS402764 |
| DK17 | Ii | PID16905 | 9/5/06 | ERS402833 |
| DK18 | 414 | PID80224 | 7/12/11 | ERS402749 |
| DK19 | 156 | PID08136 | 7/4/06 | ERS402467 |
| DK20 | 166 | PID08136 | 3/18/11 | ERS402480 |
| DK21 | 182 | PID35550 | 5/23/05 | ERS402499 |
| DK22 | 178 | PID35550 | 1/12/09 | ERS402493 |
| DK23 | 177 | PID35550 | 3/2/10 | ERS402492 |
| DK24 | 173 | PID35550 | 8/8/12 | ERS402488 |
| DK25 | 192 | PID08309 | 4/27/09 | ERS402510 |
| DK26 | 188 | PID08309 | 8/15/11 | ERS402505 |
| DK27 | 205 | PID61790 | 11/24/04 | ERS402526 |
| DK28 | 202 | PID61790 | 3/21/06 | ERS402522 |
| DK29 | 199 | PID61790 | 5/29/07 | ERS402517 |
| DK30 | 323 | PID67971 | 8/10/06 | ERS402649 |
| DK31 | 220 | PID60932 | 2/24/09 | ERS402544 |
| DK32 | 224 | PID52119 | 2/19/07 | ERS402548 |
| DK33 | 187 | PID08309 | 10/31/11 | ERS402504 |
| DK34 | 250 | PID75754 | 2/9/10 | ERS402569 |
| DK35 | 254 | PID75754 | 3/13/06 | ERS402571 |
| DK36 | 410 | PID51508 | 6/25/07 | ERS402742 |
| DK37 | 293 | PID38603 | 9/2/02 | ERS402614 |

| | | | | |
|---|---|---|---|---|
| DK38 | 349 | PID15859 | 1/16/07 | ERS402676 |
| DK39 | 355 | PID46516 | 9/27/06 | ERS402682 |
| DK40 | 381 | PID79984 | 7/29/05 | ERS402710 |
| DK41 | 366 | PID79984 | 1/8/08 | ERS402694 |
| DK42 | 371 | PID67941 | 5/30/11 | ERS402700 |
| DK43 | 373 | PID67941 | 10/15/09 | ERS402702 |
| DK44 | 336 | PID02938 | 8/11/09 | ERS402662 |
| DK45 | 396 | PID20509 | 6/6/11 | ERS402726 |
| DK46 | 324 | PID67971 | 8/13/08 | ERS402650 |
| DK47 | 339 | PID15859 | 5/18/05 | ERS402665 |
| DK48 | 340 | PID15859 | 8/24/10 | ERS402667 |
| DK49 | 341 | PID15859 | 2/7/11 | ERS402668 |
| DK50 | 386 | PID42824 | 7/8/09 | ERS402715 |
| DK51 | 388 | PID42824 | 7/24/07 | ERS402717 |
| DK52 | 406 | PID51508 | 8/9/11 | ERS402737 |
| DK53 | 434 | PID12139 | 6/28/05 | ERS402772 |

**Supplementary Table 2.** T6SS effectors known in *Pseudomonas aeruginosa*

| Effector locus | Effector name (alternative name) | T6SS | Target | Transport | References |
|---|---|---|---|---|---|
| PA0093 | Tse6 | H1 | Bacteria | PAAR | Whitney *et al.*[1] |
| PA0093 | Tas1 | H1 | Bacteria | PAAR | Ahmad *et al.*[2] |
| PA0099 | Tse7 | H1 | Bacteria | PAAR | Hachani *et al.*[3], Pissaridou *et al.*[4] |
| PA0260 | Tle3 | H2 | Bacteria | VgrG | Russell *et al.*[5], Berni *et al.*[6], Wood *et al.*[7] |
| PA0262 | VgrG2b | H2 | Trans-kingdom | VgrG | Sana *et al.*[8], Berni *et al.*[6], Wood *et al.*[7] |
| PA0807 | AmpDh3 | H2 | Bacteria | VgrG | Wang *et al.*[9] |
| PA0822 | TseV | NA | Bacteria | PAAR | Wood *et al.*[10], Wettstadt *et al.*[11], Wang *et al.*[12] |
| PA1510 | TplE (Tle4) | H2 | Trans-kingdom | VgrG | Russell *et al.*[5], Jiang *et al.*[13] |
| PA1844 | Tse1 | H1 | Bacteria | Hcp | Hood *et al.*[14], Russell *et al.*[15], Silverman *et al.*[16] |
| PA1863 | ModA | H2 | Extracellular | VgrG | Wang *et al.*[17] |
| PA2374 | TseF | H3 | Extracellular | Hcp | Lin *et al.*[18] |
| PA2684 | Tse5 (RhsP1) | H1 | Bacteria | VgrG | Hachani *et al.*[3], Whitney *et al.*[1] |
| PA2702 | Tse2 | H1 | Bacteria | Hcp | Hood *et al.*[14], Silverman *et al.*[16] |
| PA2774 | Tse4 | H1 | Bacteria | Hcp | Whitney *et al.*[1] |
| PA3290 | Tle1 | H2 | Bacteria | VgrG | Barret *et al.*[19], Russell *et al.*[5] |
| PA3484 | Tse3 | H1 | Bacteria | Hcp | Hood *et al.*[14], Russell *et al.*[15], Hachani *et al.*[20], Silverman *et al.*[16] |
| PA3487 | PldA (Tle5a) | H2 | Trans-kingdom | VgrG | Wilderman *et al.*[21], Russell *et al.*[5], Jiang *et al.*[22] |
| PA3907 | TseT | H2 | Bacteria | PAAR | Burkinshaw *et al.*[23] |
| PA4163 | Tse8 | H1 | Bacteria | VgrG | Nolan *et al.*[24] |
| PA4922 | Azu | H2 | Extracellular | VgrG | Han *et al.*[25] |
| PA5089 | PldB (Tle5b) | H2 | Trans-kingdom | VgrG | Russell *et al.*[5], Jiang *et al.*[22], Wettstadt *et al.*[11] |
| PA5265* | - | NA | NA | NA | Wettstadt *et al.*[11] |
| PA14_43100 | RhsP2 | H2 | Bacteria | PAAR | Lesic *et al.*[26], |

| | | | | | |
|---|---|---|---|---|---|
| | | | | | Hachani *et al.*[3], Jones *et al.*[27] |

*Effector has not yet been experimentally validated

**Supplementary Table 3.**

Is provided as a separate .xls document.

**Supplementary Table 4.** Bacterial strains used in this study.

| Name | Description | Source |
|---|---|---|
| WT | *Pseudomonas aeruginosa* PAO1, DSM22644 | DSMZ |
| ΔPA3487-PA3488 | PAO1 strain described above with an *in-frame* deletion of *pldA* (PA3487) and *tli5a* (PA3488) | This study |
| ΔPA0807-PA0808 | PAO1 strain described above with an *in-frame* deletion of *ampDh3* (PA0807) and (PA0808) | This study |
| ΔPA1509-PA1510 | PAO1 strain described above with an *in-frame* deletion of PA1509 and *tplE* (PA1510) | This study |
| ΔPA1844-PA1845 | PAO1 strain described above with an *in-frame* deletion of *tse1* (PA1844) and *tsi1* (PA1845) | This study |
| ΔPA2374 | PAO1 strain described above with an *in-frame* deletion of *tseF* (PA2374) | This study |
| ΔPA2684-PA2684.1 | PAO1 strain described above with an *in-frame* deletion of *tse5* (PA2684) and *tsi5* (PA2684.1) | This study |
| ΔPA2702-PA2703 | PAO1 strain described above with an *in-frame* deletion of *tse2* (PA2702) and *tsi2* (PA2703) | This study |
| ΔPA3290-PA3291 | PAO1 strain described above with an *in-frame* deletion of *tle1* (PA3290) and PA3291. | This study |
| ΔPA3484-PA3485 | PAO1 strain described above with an *in-frame* deletion of *tse3* (PA3484) and *tsi3* (PA3485) | This study |
| ΔPA3907-PA3908 | PAO1 strain described above with an *in-frame* deletion of *tseT* (PA3907) and *tsiT* (PA3908) | This study |
| ΔPA4163-PA4164 | PAO1 strain described above with an *in-frame* deletion of *tse8* (PA4163) and *tsi8* (PA4164) | This study |
| ΔPA4922 | PAO1 strain described above with an *in-frame* deletion of *azu* (PA4922) | This study |
| ΔPA5086-PA5089 | PAO1 strain described above with an *in-frame* deletion of *pldB* (PA5089) and PA5086-PA5088 | This study |
| WT YFP | PAO1 strain described above constitutively expressing sYFP2 after Tn7-143[28] had been inserted in the chromosome and the plasmid eliminated. | This study |
| ΔPA3487-PA3488 YFP | PAO1 deletion strain described above constitutively expressing sYFP2 after Tn7-143[28] had been inserted in the chromosome and the plasmid eliminated. | This study |
| ΔPA3907-PA3908 YFP | PAO1 deletion strain described above constitutively expressing sYFP2 after Tn7-143[28] had been inserted in the chromosome and the plasmid eliminated. | This study |

**Supplementary Table 5.** Oligonucleotide primers for cloning used in this study.

| Name | Primer (5’-3’; restriction sites underlined) |
|---|---|
| P1_ PA3487-PA3488 | AAGCTTCGTGGAGGCGAACAGCTACAGCG GTTCAAGG |
| P2_ PA3487-PA3488 | CGATGTGGGTTTCAGCGCGCCAGCCGCTTCTTCTGC<br>AACATGGATCAGTCC |
| P3_ PA3487-PA3488 | GGACTGATCCATGTTGCAGAAGAAGCGGCTGGCGCG<br>CTGAAACCCACATCG |
| P4_ PA3487-PA3488 | TCTAGAGGTCTGCGGCAACAGGGCGTTGACCTGCTCGGCG |
| P1_ PA0807-PA0808 | AAGCTTACCCCGACCTGTTCCCATCCTTC |
| P2_ PA0807-PA0808 | CAATGCTCAGCCATCGAGACCGTCGATGGTCAGCATGGTTTTC |
| P3_ PA0807-PA0808 | GAAAACCATGCTGACCATCGACGGTCTCGATGGCTGAGCATTG |
| P4_ PA0807-PA0808 | GGTACCGGCGAGGTGCTCCAAGGGTTC |
| P1_ PA1509-PA1510 | AAGCTTCGATCAGTTACCACGGACATGCAACACCTAGG |
| P2_ PA1509-PA1510 | CACATGAGCAGCGAACCGCTGGACGACCTATAGGAGC |
| P3_ PA1509-PA1510 | GCTCCTATAGGTCGTCCAG CGGTTCGCTGCTCATGTG |
| P4_ PA1509-PA1510 | GGTACCGAGGCGTCGTGGTGCAGGTG |
| P1_ PA1844-PA1845 | AAGCTTGGTGTCGAAGATGGTCTCGATCAGGATCAGGTCG |
| P2_ PA1844-PA1845 | CCCCATGAAACTGCTCGCCCCCAGGGCCAGTTGATTCAG |
| P3_ PA1844-PA1845 | CTGAATCAACTGGCCCTGGG GGCGAGCAGTTTCATGGGG |
| P4_ PA1844-PA1845 | GGTACC CAAAGTGGCGTATTCGTCGTCGCTCAGC |
| P1_ PA2374 | TCTAGACCAGGAAGTGATCGTCACCTTCATCGACGG |
| P2_ PA2374 | GCGCCTAGGGCTCCGCCAGGCCGGATGCCGCCATGCGC |
| P3_ PA2374 | GCGCATGGCGGCATCCGGCCTGGCGGAGCCCTAGGCGC |
| P4_ PA2374 | GGTACCGCACCTCGTCGAAGCTGCCCG |
| P1_ PA2684-PA2684.1 | GAATTCCGCCAGGGCACAGC |
| P2_ PA2684-PA2684.1 | GGCCTCACTCGTCCAGGTCGGTAGGCCGCTCATCTAT |
| P3_ PA2684-PA2684.1 | ATAGATGAGCGGCCTACCGACCTGGACGAGTGAGGCC |
| P4_ PA2684-PA2684.1 | GGTACCGAGCAGACGGTGC |
| P1_ PA2702-PA2703 | GGTACCTCG CCG TGC TCT CCA GCC TG |
| P2_ PA2702-PA2703 | GCCGTCAGGATGCCGGCTCGTAGTCGTAGGACATGCGTG |
| P3_ PA2702-PA2703 | CACGCATGTCCTACGACTAC GAGCCGGCATCCTGACGGC |
| P4_ PA2702-PA2703 | TCTAGAGGTTTCGCCTGCGAGTTGCACTTCG |
| P1_ PA3290-PA3291 | AAGCTTGCAGGTGTCAGCCTGGATGC |
| P2_ PA3290-PA3291 | GGGTATGTCATGCAGAAGTTGAT<br>GATGGAATCGCTTTGAACCTCGAC |
| P3_ PA3290-PA3291 | GTCGAGGTTCAAAGCGATTCCAT<br>CATCAACTTCTGCATGACATACCC |
| P4_ PA3290-PA3291 | GGTACCTGAAAGCTGTCGTGGAATGGGAGAAAGATCC |
| P1_ PA3484-PA3485 | TCTAGAGAGGGCTCGATAGAATGAACGCATGGAAGC |
| P2_ PA3484-PA3485 | GGTTCTATTGCAACCTGGCGCTGGTGGCGGTCATCGGTC |
| P3_ PA3484-PA3485 | GACCGATGACCGCCACCAGC GCCAGGTTGCAATAGAACC |
| P4_ PA3484-PA3485 | GGTACCAATGCAGGTCGCTTTCGTCGTACTGC |

| | |
|---|---|
| P1_ PA3907-PA3908 | AAGCTTCGGGAGCGAGCTGGACGAAATCG |
| P2_ PA3907-PA3908 | GTCGTCAGCCGCGCAGCGGGCCTATGCCGCTCATGCCG |
| P3_ PA3907-PA3908 | CGGCATGAGCGGCATAGGCCCGCTGCGCGGCTGACGAC |
| P4_ PA3907-PA3908 | TCTAGACCTGACCATCGCGGTGAACCACCTGAAGTCG |
| P1_ PA4163-PA4164 | AAGCTTCAGGTGCGAGTCACCCCGACCAG |
| P2_ PA4163-PA4164 | GCGCTCAGTCGCGCAGGGCGGTGACCTCGATCATGCTG |
| P3_ PA4163-PA4164 | CAGCATGATCGAGGTCACC GCCCTGCGCGACTGAGCGC |
| P4_ PA4163-PA4164 | GGTACCCGATCCTCGCCCAGCCG |
| P1_ PA4922 | AAGCTTGAATGTGCCTGTCGCAACCAGGTCTTCAAGTC |
| P2_ PA4922 | CTCCATGCTACGTAAACTCCTGACCCTGAAGTGATGCG |
| P3_ PA4922 | CGCATCACTTCAGGGTCAG GAGTTTACGTAGCATGGAG |
| P4_ PA4922 | GGTACCGTCGTAGAAACCGTTCACTTCGAGCAGGC |
| P1_ PA5086-PA5089 | AAGCTTGGATGACAGCGGTAGGACAATGAATAAAGATGGC |
| P2_ PA5086-PA5089 | CGCCATGAGCGACCTGTACAAGCCGGAGCGGTAAGTAC |
| P3_ PA5086-PA5089 | GTACTTACCGCTCCGGCTT GTACAGGTCGCTCATGGCG |
| P4_ PA5086-PA5089 | GGTACCGGTTTCCTGGAGGGCGATCCG |

75

**Supplementary Table 6.** Diverse *P. aeruginosa* isolate collection from Hilker *et al.*[29]

| Clone | Isolate name | Sequence Read Archive (SRA) accession number |
|---|---|---|
| C40A | NN2 | ERR371809 |
| D421 | RN3 | ERR371810 |
| F469 | 60P57PA | ERR371811 |
| 0C2E | RP1 | ERR371812 |
| E429 | 15108/-1 | ERR371813 |
| 239A | 13121/-1 | ERR371814 |
| 2C22 | 57P31PA | ERR371815 |
| F429 | A5803 | ERR371816 |
| B420 | 120SD3 | ERR371817 |
| EA0A | 39177 | ERR371818 |
| 812 | 27103 | ERR371819 |
| 2C1A | 18P17PA | ERR371820 |
| 1BAE | KK1 | ERR371821 |
| 3C2A | TR1 | ERR371822 |
| EC2A | PT22 | ERR371823 |
| EC21 | 100 | ERR371824 |
| 843A | BP35 | ERR371825 |
| 822 | E501 | ERR371826 |
| 149A | 120SB2 | ERR371827 |
| 478A | MA41A.1 | ERR371828 |

**Supplementary Table 7.** Available *P. aeruginosa* complete genomes on NCBI as of December 2020.

| Strain | Sequence Accession |
|---|---|
| PAO1 | AE004091 |
| NCGM2.S1 | AP012280 |
| NCGM1900 | AP014622 |
| NCGM1984 | AP014646 |
| NCGM257 | AP014651 |
| 8380 | AP014839 |
| IOMTU 133 | AP017302 |
| UCBPP-PA14 | CP000438 |
| PA7 | CP000744 |
| M18 | CP002496 |
| DK2 | CP003149 |
| PA1 | CP004054 |
| PA1R | CP004055 |
| B136-33 | CP004061 |
| RP73 | CP006245 |
| MTB-1 | CP006853 |
| SCV20265 | CP006931 |
| LES431 | CP006937 |
| YL84 | CP007147 |
| F22031 | CP007399 |
| VRFPA04 | CP008739 |
| F23197 | CP008856 |
| isolate F30658 | CP008857 |
| F63912 | CP008858 |
| H5708 | CP008859 |
| H27930 | CP008860 |
| isolate H47921 | CP008861 |
| M1608 | CP008862 |
| isolate M37351 | CP008863 |
| W60856 | CP008864 |
| S86968 | CP008865 |
| T38079 | CP008866 |
| isolate T52373 | CP008867 |
| isolate T63266 | CP008868 |

| | |
|---|---|
| W16407 | CP008869 |
| W36662 | CP008870 |
| W45909 | CP008871 |
| X78812 | CP008872 |
| isolate F9670 | CP008873 |
| FRD1 | CP010555 |
| Carb01 63 | CP011317 |
| ATCC 27853 | CP011857 |
| DSM 50071 | CP012001 |
| F9676 | CP012066 |
| PA_D2 | CP012578 |
| PA_D5 | CP012579 |
| PA_D9 | CP012580 |
| PA_D16 | CP012581 |
| PA_D21 | CP012582 |
| PA_D22 | CP012583 |
| PA_D25 | CP012584 |
| PA_D1 | CP012585 |
| PA1RG | CP012679 |
| N15-01092 | CP012901 |
| PAER4_119 | CP013113 |
| VA-134 | CP013245 |
| SCVFeb | CP013477 |
| SCVJan | CP013478 |
| NHmuc | CP013479 |
| 12-4-4(59) | CP013696 |
| USDA-ARS-USMARC-41639 | CP013989 |
| DH01 | CP013993 |
| PA_154197 | CP014866 |
| N17-1 | CP014948 |
| PA1088 | CP015001 |
| PA8281 | CP015002 |
| PA11803 | CP015003 |
| ATCC 27853 | CP015117 |
| BAMC 07-48 | CP015377 |
| PA121617 | CP016215 |
| RIVM-EMC2982 | CP016955 |
| DN1 | CP017099 |
| ATCC 15692 | CP017149 |

| | |
|---|---|
| PA83 | CP017293 |
| PA_150577 | CP017306 |
| FA-HZ1 | CP017353 |
| isolate B10W | CP017969 |
| L10 | CP019338 |
| CR1 | CP020560 |
| E6130952 | CP020603 |
| PAK | CP020659 |
| PASGNDM345 | CP020703 |
| PASGNDM699 | CP020704 |
| Pa124 | CP021774 |
| Pa58 | CP021775 |
| Pa84 | CP021999 |
| Pa127 | CP022000 |
| Pa1207 | CP022001 |
| Pa1242 | CP022002 |
| LW | CP022478 |
| Ocean-1175 | CP022525 |
| Ocean-1155 | CP022526 |
| CCUG 70744 | CP023255 |
| PPF-1 | CP023316 |
| 12939 | CP024477 |
| PA59 | CP024630 |
| PB369 | CP025049 |
| PB368 | CP025050 |
| PB353 | CP025051 |
| PB354 | CP025053 |
| PB350 | CP025055 |
| PB367 | CP025056 |
| F5677 | CP026680 |
| AR_0360 | CP027165 |
| AR_0357 | CP027166 |
| AR_0356 | CP027169 |
| AR_0354 | CP027171 |
| AR_0353 | CP027172 |
| AR_0230 | CP027175 |
| AR_0095 | CP027538 |
| YB01 | CP028132 |
| MRSN12280 | CP028162 |

| | |
|---|---|
| WCHPA075019 | CP028584 |
| IMP67 | CP028848 |
| IMP68 | CP028849 |
| JB2 | CP028917 |
| IMP66 | CP028959 |
| AR445 | CP029088 |
| AR444 | CP029089 |
| AR442 | CP029090 |
| AR441 | CP029093 |
| AR439 | CP029095 |
| 24Pae112 | CP029605 |
| AR_0446 | CP029660 |
| K34-7 | CP029707 |
| AR_0110 | CP029745 |
| AR_458 | CP030327 |
| AR_455 | CP030328 |
| AR_460 | CP030351 |
| HS9 | CP030861 |
| Y31 | CP030910 |
| Y71 | CP030911 |
| Y82 | CP030912 |
| Y89 | CP030913 |
| 97 | CP031449 |
| PABL012 | CP031659 |
| PABL017 | CP031660 |
| E80 | CP031677 |
| PAO1161 | CP032126 |
| AR_0111 | CP032256 |
| PA34 | CP032552 |
| BA7823 | CP032569 |
| 268 | CP032761 |
| BA15561 | CP033432 |
| SP4528 | CP033439 |
| H26027 | CP033684 |
| H26023 | CP033685 |
| H25883 | CP033686 |
| FDAARGOS_532 | CP033771 |
| FDAARGOS_505 | CP033832 |
| FDAARGOS_571 | CP033833 |

| | |
|---|---|
| FDAARGOS_570 | CP033835 |
| FDAARGOS_501 | CP033843 |
| IMP-13 | CP034354 |
| B41226 | CP034368 |
| SP4371 | CP034369 |
| SP4527 | CP034409 |
| GIMC5015:PAKB6 | CP034429 |
| SP2230 | CP034434 |
| B14130 | CP034435 |
| B17932 | CP034436 |
| AES1M | CP037925 |
| AES1R | CP037926 |
| PABL048 | CP039293 |
| T2436 | CP039989 |
| T2101 | CP039990 |
| PA298 | CP040126 |
| C79 | CP040684 |
| FDAARGOS_767 | CP041008 |
| FDAARGOS_610 | CP041013 |
| AZPAE15042 | CP041354 |
| A681 | CP041771 |
| 243931 | CP041772 |
| 519119 | CP041773 |
| 60503 | CP041774 |
| ST773 | CP041945 |
| HOU1 | CP042268 |
| CCUG 51971 | CP043328 |
| E90 | CP044006 |
| AG1 | CP045739 |
| CFSAN084950 | CP045768 |
| CF39S | CP045916 |
| 1811-18R001 | CP046060 |
| 1811-13R031 | CP046061 |
| KRP1 | CP046069 |
| INP-43 | CP047592 |
| RD1-3 | CP047697 |
| LESB58 | FM209186 |
| NCTC10332; NCTC10332 | LN831024 |
| DK1 substr. NH57388A | LN870292 |

| | |
|---|---|
| PAO1_Orsay | LN871187 |
| isolate paerg002 | LR130527 |
| isolate paerg000 | LR130528 |
| isolate paerg003 | LR130530 |
| isolate paerg004 | LR130531 |
| isolate paerg009 | LR130533 |
| isolate paerg005 | LR130534 |
| isolate paerg011 | LR130535 |
| isolate paerg010 | LR130536 |
| isolate paerg012 | LR130537 |
| NCTC11445 | LR134308 |
| NCTC12903 | LR134309 |
| NCTC13715 | LR134330 |
| NCTC10728 | LR134342 |
| NCTC13620 | LR590472 |
| NCTC13359 | LR590473 |
| NCTC13618 | LR590474 |
| PAK | LR657304 |
| isolate 1 | LS998783 |
| isolate PA14Or_reads | LT608330 |
| isolate PcyII-10 | LT673656 |
| isolate early isolate NN2 (clone C) | LT883143 |
| isolate RW109 | LT969520 |

### References


1. Whitney, J. C. *et al.* Genetically distinct pathways guide effector export through the type VI secretion system. *Mol. Microbiol.* **92**, 529–542 (2014).
2. Ahmad, S. *et al.* An interbacterial toxin inhibits target cell growth by synthesizing (p)ppApp. *Nature* **575**, 674–678 (2019).
3. Hachani, A., Allsopp, L. P., Oduko, Y. & Filloux, A. The VgrG proteins are 'à la carte' delivery systems for bacterial type VI effectors. *J. Biol. Chem.* **289**, 17872–17884 (2014).
4. Pissaridou, P. *et al.* The Pseudomonas aeruginosa T6SS-VgrG1b spike is topped by a PAAR protein eliciting DNA damage to bacterial competitors. *Proc. Natl. Acad. Sci. U. S. A.* **115**, 12519–12524 (2018).
5. Russell, A. B. *et al.* Diverse type VI secretion phospholipases are functionally plastic antibacterial effectors. *Nature* **496**, 508–512 (2013).
6. Berni, B., Soscia, C., Djermoun, S., Ize, B. & Bleves, S. A type VI secretion system trans-kingdom effector is required for the delivery of a novel antibacterial toxin in Pseudomonas aeruginosa. *Front. Microbiol.* **10**, (2019).
7. Wood, T. E. *et al.* The Pseudomonas aeruginosa T6SS Delivers a Periplasmic Toxin that Disrupts Bacterial Cell Morphology. *Cell Rep.* **29**, 187-201.e7 (2019).
8. Sana, T. G. *et al.* Internalization of Pseudomonas aeruginosa strain PAO1 into epithelial cells is promoted by interaction of a T6SS effector with the microtubule network. *MBio* **6**, 1–11 (2015).
9. Wang, T. *et al.* A type VI secretion system delivers a cell wall amidase to target bacterial competitors. *Mol. Microbiol.* **114**, 308–321 (2020).
10. Wood, T. E., Howard, S. A., Wettstadt, S. & Filloux, A. PAAR proteins act as the 'sorting hat' of the type VI vsecretion system. *Microbiol. (United Kingdom)* **165**, 1208–1218 (2019).
11. Wettstadt, S., Wood, T. E., Fecht, S. & Filloux, A. Delivery of the Pseudomonas aeruginosa phospholipase effectors PldA and PldB in a VgrG- And H2-T6SS-dependent manner. *Front. Microbiol.* **10**, 1–18 (2019).
12. Wang, S., Geng, Z., Zhang, H., She, Z. & Dong, Y. The Pseudomonas aeruginosa PAAR2 cluster encodes a putative VRR-NUC domain-containing effector. *FEBS J.* **288**, 5755–5767 (2021).
13. Jiang, F. *et al.* The Pseudomonas aeruginosa Type VI Secretion PGAP1-like Effector Induces Host Autophagy by Activating Endoplasmic Reticulum Stress. *Cell Rep.* **16**, 1502–1509 (2016).
14. Hood, R. D. *et al.* A Type VI Secretion System of Pseudomonas aeruginosa Targets a Toxin to Bacteria. *Cell Host Microbe* **7**, 25–37 (2010).
15. Russell, A. B. *et al.* Type VI secretion delivers bacteriolytic effectors to target cells. *Nature* **475**, 343–347 (2011).
16. Silverman, J. M. *et al.* Haemolysin Coregulated Protein Is an Exported Receptor and Chaperone of Type VI Secretion Substrates. *Mol. Cell* **51**, 584–593 (2013).
17. Wang, T. *et al.* Pseudomonas aeruginosa T6SS-mediated molybdate transport contributes to bacterial competition during anaerobiosis. *Cell Rep.* **35**, 108957 (2021).
18. Lin, J. *et al.* A Pseudomonas T6SS effector recruits PQS-containing outer membrane vesicles for iron acquisition. *Nat. Commun.* **8**, 1–12 (2017).
19. Barret, M., Egan, F., Fargier, E., Morrissey, J. P. & O'Gara, F. Genomic analysis of the

type VI secretion systems in Pseudomonas spp.: Novel clusters and putative effectors uncovered. *Microbiology* **157**, 1726–1739 (2011).

20. Hachani, A. *et al.* Type VI secretion system in Pseudomonas aeruginosa: Secretion and multimerization of VgrG proteins. *J. Biol. Chem.* **286**, 12317–12327 (2011).
21. Wilderman, P. J., Vasil, A. I., Johnson, Z. & Vasil, M. L. Genetic and biochemical analyses of a eukaryotic-like phospholipase D of Pseudomonas aeruginosa suggest horizontal acquisition and a role for persistence in a chronic pulmonary infection model. *Mol. Microbiol.* **39**, 291–304 (2001).
22. Jiang, F., Waterfield, N. R., Yang, J., Yang, G. & Jin, Q. A Pseudomonas aeruginosa type VI secretion phospholipase D effector targets both prokaryotic and eukaryotic cells. *Cell Host Microbe* **15**, 600–610 (2014).
23. Burkinshaw, B. J. *et al.* A type VI secretion system effector delivery mechanism dependent on PAAR and a chaperone-co-chaperone complex. *Nat. Microbiol.* **3**, 632–640 (2018).
24. Nolan, L. M. *et al.* Identification of Tse8 as a Type VI secretion system toxin from Pseudomonas aeruginosa that targets the bacterial transamidosome to inhibit protein synthesis in prey cells. *Nat. Microbiol.* **6**, 1199–1210 (2021).
25. Han, Y. *et al.* A Pseudomonas aeruginosa type VI secretion system regulated by CueR facilitates copper acquisition. *PLoS Pathog.* **15**, e1008198 (2019).
26. Lesic, B., Starkey, M., He, J., Hazan, R. & Rahme, L. G. Quorum sensing differentially regulates Pseudomonas aeruginosa type VI secretion locus I and homologous loci II and III, which are required for pathogenesis. *Microbiology* **155**, 2845–2855 (2009).
27. Jones, C., Hachani, A., Manoli, E. & Filloux, A. An rhs gene linked to the second type VI secretion cluster is a feature of the pseudomonas aeruginosa strain PA14. *J. Bacteriol.* **196**, 800–810 (2014).
28. Schlechter, R. O. *et al.* Chromatic Bacteria – A Broad Host-Range Plasmid and Chromosomal Insertion Toolbox for Fluorescent Protein Expression in Bacteria. *Front. Microbiol.* **9**, (2018).
29. Hilker, R. *et al.* Interclonal gradient of virulence in the Pseudomonas aeruginosa pangenome from disease and environment. *Environ. Microbiol.* **17**, 29–46 (2015).
